# Dynamic and Structural Modeling of the Specificity in Protein-DNA Interactions Guided by Binding Assay and Structure Data

**DOI:** 10.1101/288795

**Authors:** Cheng Tan, Shoji Takada

**Affiliations:** Department of Biophysics, Graduate School of Science, Kyoto University, Kyoto 606-8502, Japan

**Keywords:** PWMcos, protein-DNA sequence-specific binding, transcription factor target search

## Abstract

How transcription factors (TFs) recognize their DNA sequences is often investigated complementarily by high-throughput protein binding assays and by structural biology experiments. The former quantifies the specificity of TF binding sites for numerous DNA sequences, often represented as the position-weight-matrix (PWM). The latter provides mechanistic insights into the interactions via the protein-DNA complex structures. However, these two types of data are not readily integrated. Here, we propose and test a new modeling method that incorporates the PWM with complex structure data. Based on pre-tuned coarse-grained models for proteins and DNAs, we model the specific protein-DNA interactions, PWMcos, in terms of an orientation-dependent potential function, which enables us to perform molecular dynamics simulations at unprecedentedly large scales. We show that the PWMcos model reproduces subtle specificity in the protein-DNA recognition. During the target search in genomic sequences, TF moves on highly rugged landscapes and occasionally flips on DNA depending on the sequence. The TATA-binding protein exhibits two remarkably distinct binding modes, of which frequencies differ between TATA-containing and TATA-less promoters. The PWMcos is general and can be applied to any protein-DNA interactions given their PWMs and complex structure data are available.

## INTRODUCTION

Specificity in protein-DNA interactions plays central roles in many molecular genetic processes.^1–3^ Transcription factors (TFs), for example, search and recognize their target DNA sequences via specific protein-DNA interactions. Recent high-throughput technologies such as the systematic evolution of ligands by exponential enrichment (SELEX)^4^ and the protein-binding microarrays (PBMs)^5^ have greatly expanded our knowledge of the sequence preferences at TF binding sites, which by now became large-scale datasets of relative binding affinities, i.e., specificities for thousands of TFs.^6^ Theoretical models have been developed to represent the specificity in tractable forms.^7^ Among them, the simplest and most popular one is the position weight matrix (PWM). With the primary assumption that each position in the protein binding site contributes independently to recognition, the model assigns a score to each possible base (among A, C, G, and T) in each position, the sum of which gives a total score that can be used as an estimation of specificity.^7,8^ Many different methods have been developed to assign the elements in the PWM, such as the probabilistic modeling^9^ and the energy modeling.^10^ The PWM can be translated into graphical representations (e.g., the “sequence logo”), which are nowadays extensively used in literature.^11^ In PWM representations, however, the DNA sequences in the TF binding sites play central roles, while TFs themselves are treated implicitly. For example, crosstalk among TFs is only inferred from the correlation in PWMs. How bound TFs operate on transcription machinery, for example, is hardly addressed with the DNA sequence-centric view.

In this sense, complementary insights on the TF recognition of its DNA sequence and TF operations on transcription can be gained via structural biology experiments, where X-ray crystallographic structures of TF-DNA complexes provide mechanistic understanding on the recognition and the operation. TFs can directly recognize the unique chemical features of nucleobases from both the major and minor grooves; they also can read out the sequence-dependent geometric characteristics of DNA.^12^ Crosstalk among TFs and with nucleosomes as well as the action of TFs on transcription machinery can directly be represented as physical interactions. However, structural biology is relatively time-consuming and cannot easily be performed high-throughput so that the structure can be obtained only for, at most, a handful of DNA sequences per TF.

While high-throughput protein binding assays and structural biology experiments are clearly complementary, it is not straightforward to systematically integrate the two types of data. The purpose of this study is to integrate the PWM and the complex structure data into a new method of molecular dynamics (MD) simulations. Using MD simulations, one can investigate not only structural but also dynamical aspects in the TF recognition of DNA sequences and TF operations on transcription activation and repression. These processes are not easily observed experimentally because of limitation in both spatial and temporal resolutions and thus computer modeling is highly desired.

MD simulations are potentially powerful to investigate dynamic protein-DNA interactions because of their high resolutions in space and time. Yet, fully atomistic MD simulations are currently limited to shorter timescales than necessary. Recently, due to the development of well-tuned modeling, coarse-grained (CG) molecular simulations become popular providing many promising results on TF dynamics as well as chromatin dynamics.^13–20^ Among many possible resolutions in CG modeling, here we take one in which each amino acid is simplified as one particle and each nucleotide is represented by three particles, each for sugar, phosphate, and base. For proteins, atomic-interaction based coarse-grained (AICG2+) model captures local structural dynamics of protein around the native structure accurately and has been applied to various systems.^21–23^ For DNA, by introducing the sequence-dependent structural and mechanical properties of DNA, the latest 3-site DNA model (3SPN.2C) can be used to model the indirect readout of DNA sequence by protein.^24–27^ New methods of re-calculating protein charge distributions have also been developed to improve the accuracy of modeling electrostatic interactions between protein and DNA.^28,29^ On the other hand, specific interactions between proteins and DNAs are only limited to a rather crude model, in which we assume the perfect specificity to one site in the background of no specific interaction at all.^30,31^ This all-or-none type modeling cannot give a proper prediction on cases where more than one consensus pieces or pseudo binding sites exist. Given these, here we develop a generic specificity model for protein-DNA interaction based both on PWMs and on complex structure data.

One clear example that is benefitted by this MD method is the TF search process. Due to the relatively short length of DNA sequences recognized by TFs, especially those in eukaryotes, there are numerous on- and off-targets for TFs to bind, which undoubtedly affects the target searching dynamics of proteins. Pseudo-specific sequences exist more frequently than the target, which can make the TF landscape on genomic DNA highly rugged and the search dynamics highly non-trivial. With the method developed here, one can easily perform TF search dynamic simulations on genomic DNA sequences, addressing structural dynamics therein.

In this paper, we present a new model of sequence-specific protein-DNA interactions for CG MD simulations. Our basic idea is to construct a new potential that incorporates both the PWM data and the *co*mplex *s*tructural information (Figure 1), based on which we termed the method as PWMcos. With the assumption of additivity of independent mononucleotide contributions,^7^ the elements in PWM were properly translated into binding energies. These energies were then introduced into MD simulations by decorating them with carefully designed structure-based modulating functions. By applying this new model to two DNA-binding proteins, PU.1 and the TATA-binding protein (TBP), we demonstrated the power of the model to capture the thermodynamic and kinetic properties of the protein-DNA binding. Our model reproduced the relative binding affinities of the simulated proteins on the consensus sequence and pseudo targets. Our results also predicted a DNA sequence-dependent flipping of PU.1, and a transition between two markedly distinct binding modes of TBP during the target searching process.

**Figure 1.**
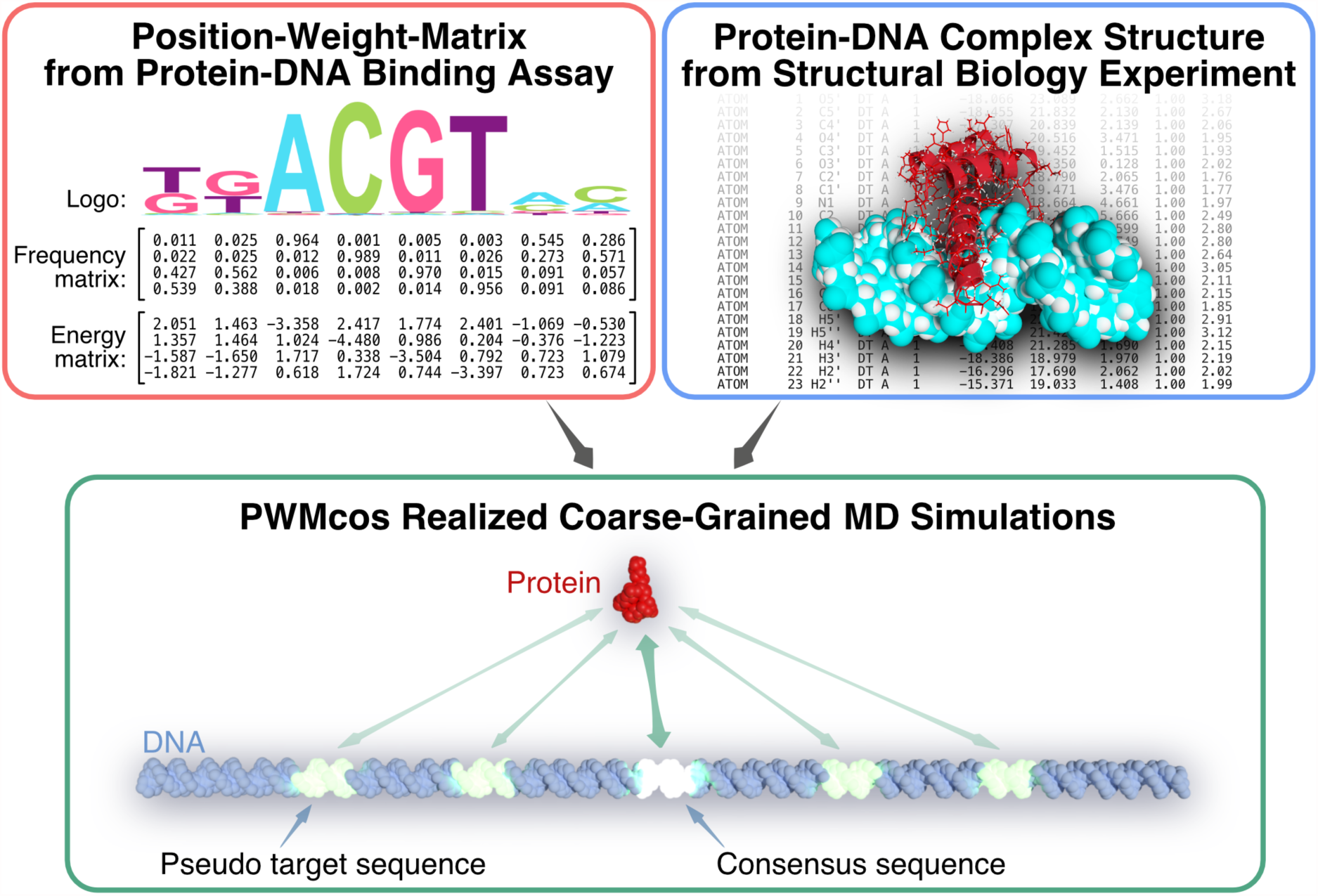
The PWMcos models the sequence-specific protein-DNA interactions guided by the position-weight-matrix (PWM) and the complex structure data. High-throughput protein-DNA binding assays provide the PWM and the “sequence logo” (top-left). Atomic structures of protein-DNA complexes provide mechanistic interaction information (top-right). The PWMcos that integrate these two types of data to build a new sequence-specific protein-DNA interaction potential can be used for coarse-grained MD simulations (bottom).

## METHODS

### Coarse-grained models for proteins and DNAs

First, we briefly describe the CG models developed for proteins and DNAs. For the CG model of proteins, we used the AICG2+ model,^21^ in which each amino acid is represented by a CG particle located at the *C*_α_ atom (see Supporting Information for more details). With a consistent resolution of roughly ten heavy atoms per one CG particle, we used the 3SPN.2C model for DNA,^24,25^ with three CG particles per one nucleotide, each representing phosphate (P), sugar (S), and base (B). Notably, one of the advanced features of the 3SPN.2C DNA model is its ability to mimic the sequence-dependent structural and mechanical properties of duplex DNA, which provides a useful framework for the “indirect readouts” in protein-DNA recognition.^25^

The carefully tuned combination of these two models has gained success in simulating protein-DNA sequence-nonspecific binding, where only electrostatic and excluded-volume effects were considered between proteins and DNAs.^26,27^ As the dominant part of the sequence-nonspecific interactions between protein and DNA, electrostatics is modeled by the Debye-Hückel theory, with charges on the *C*_α_ particles parameterized by the RESPAC method.^28^ Note that the charges of the phosphate groups were kept −0.6*e* when calculating intra-DNA interactions,^24^ whereas for protein-DNA interactions we used the value of −1.0*e*, as suggested by previous works.^26,27^ In this study, we set the ionic strength to be 150mM. As for the excluded volume interactions, we used a set of residue-type dependent radii based on statistics over protein-DNA complex structures.^27^

On top of these models, we introduce the PWMcos energy functions into CG MD simulations in the next section.

### Modeling Sequence Specific Protein-DNA Interactions

In the present work, there are two prerequisites for the CG modeling of the sequence-specific protein-DNA interactions; the PWM and the protein-DNA complex structure data. Given these data, the PWMcos model can provide energy and force formula that reproduce the DNA sequence specificity in MD simulations.

We assume that the physical interactions can be modeled based on the experimentally provided complex structure. For a given complex structure of a TF with its recognition element, we define that an amino acid in the TF and a base in the DNA is in a “native contact” when at least one of heavy-atoms in the amino acid is within 4Å from one of the base atoms. We list all the native contact pairs, the *m*-th bases *B*_*m*_ and the *j*-th amino acids, together with their local geometric factors defined below (termed the *m*, *j*-type interaction). For DNAs, the index *m* goes from 1 to *M*. *M*, with an order of 10 or less, is the total number of base pairs involved in the interface to the TF in the complex structure (Figure 2A).

**Figure 2.**
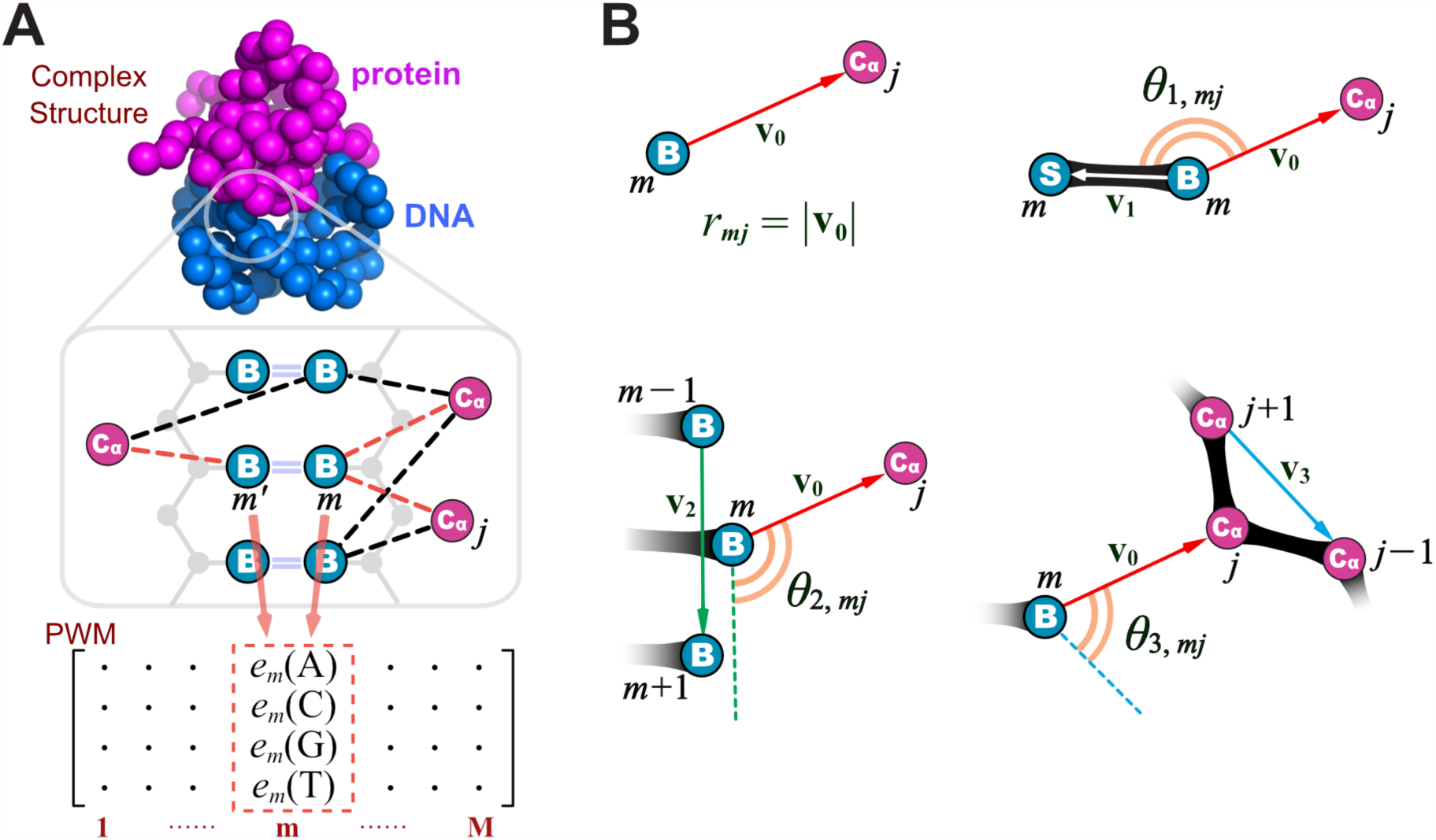
Representation of interactions between coarse-grained protein and DNA models. (A) Upper: the complex structure of a protein binding to its consensus DNA sequence. Middle: Cartoon for the native contacts. We define a native contact (dashed lines) between a base *B* (blue) and an amino acid *C*_α_ (purple). Note that one base *B*_*m*_ can form contacts with multiple *C*_α_s. The base that makes Watson-Crick base pair with *B*_*m*_ is denoted as *B*_*m*′_. DNA backbone phosphates and sugars are shown in grey. Native contacts formed between *B*_*m*_-*B*_*m*<_ and protein residues are in red. Lower: the *m*-th column in the PWM represents the recognition specificity that are contributed by all the contacts formed with the base pair *B*_*m*_-*B*_*m*′_. (B) Definition of the distance and angles involved in *U*_*mj*_ calculation. Top left: *r*_*mj*_ is the length of vector **v**_0_ connecting *B*_*m*_ and *C*_*αj*_; top right: *θ*_1,*mj*_ is the angle between vectors **v**_0_ and **v**_1_ connecting *B*_*m*_ and *S*_*m*_; bottom left: *θ*_2,*mj*_ is the angle between vectors **v**_0_ and **v**_2_ connecting *B*_*m*−1_ and *B*_*m*+1_; bottom right: *θ*_3,*mj*_ is the angle between vectors **v**_0_ and **v**_3_ connecting *C*_*α*(*j*+1)_ and *C*_*α*(*j*−1)_.

The sequence specificity can be modeled by the PWM. We denote the PWM element as *e*_*m*_(*b*), where 1 ≤ *m* ≤ *M* is the index for base and *b* ∈ {*A*, *C*, *G*, *T*} represents the base type. This index *m* should coincide with that in the native contact list described above (often, the length of PWM in the database is longer than the total number of bases appeared in the complex interface. In this case, we trim the PWM length). This index *m* should not be confused with the index *i* used below for the DNA bases in the simulation system; the simulation system contains typically much more bases. We denote the base index in the complementary strand that forms the base pair with *m*-th bases as *m*′. In the PDB structure, often both *B*_*m*_ and *B*_*m*′_ form native contacts with protein amino acids. In this case, we consider both bases contribute to the same PWM element *e*_*m*_(*b*) (Figure 2A).

Given these, we write the sequence-specific PWMcos energy (*E*_PWMcos_) as:

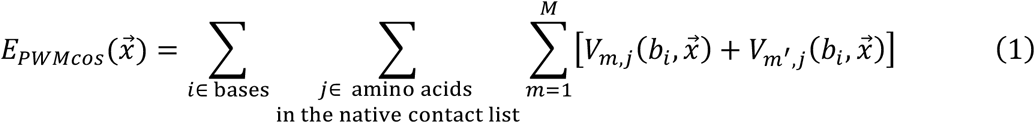

Here, 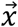 collectively represents coordinates of all the particles. 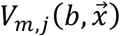 is the potential energy function that describes interactions between the *m*-th base and the *j*-th amino acids in the native complex (i.e., the *m*, *j*-type interaction) when the base type in the *m*-th base is *b* (this base type *b* may be different from that in the PDB structure). Then, the term 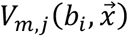 represents that the *i*-th DNA base in the simulation system interacts with the *j*-th amino acids using the *m*, *j*-type interaction. Notably, in practice, many of 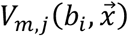 are zero if the distance between the *i*-th base and the *j*-th amino acid is larger than the characteristic distance between the *m*-th base and the *j*-th amino acids in the *m*, *j*-type interaction at the PDB structure.

To incorporate information from both the PWM and the complex structure data, we defined 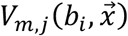 as a product of two terms:

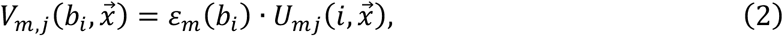

where *ε*_*m*_(*b*) is determined from the PWM and 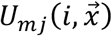 is a coordinate-dependent modulating function for the *m*, *j*-type interaction, which works for the *i*, *j* pair in the simulation system. *ε*_*m*_(*b*) is defined as the following:

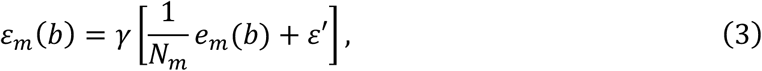

where *N*_*m*_ is the total number of native contacts formed by the base pair *B*_*m*_-*B*_*m*′_ with any amino acid. Note that for *B*_*m*′_ we have 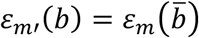, where 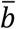 is the complementary base of *b*. In the current work, we computed *e*_*m*_(*b*) by:

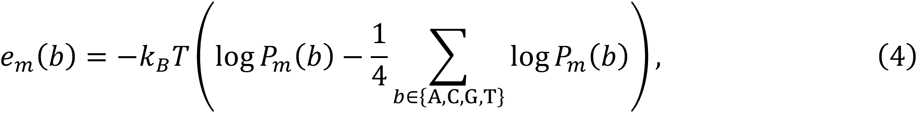

where *P*_*m*_(*b*) is the probability of finding base type *b* at the binding position *m*, *k*_*B*_ is the Boltzmann constant and *T* is temperature. Note that, in principle, *e*_*m*_ could be defined by other PWM modeling methods.^4,7,10,32,33^ However, the frequency matrix (for computing *P*_*m*_(*b*)) is the most common form of PWM in popular databases and publications. Additionally, *γ* and *ε*′ in Equation (3) are two parameters that can be calibrated by matching simulated thermodynamic quantities, such as binding affinity, to the experimental results (see Supporting Information for more details about the calibration).

The second term in Equation (2), 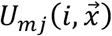, is defined as a product of four single variable functions:

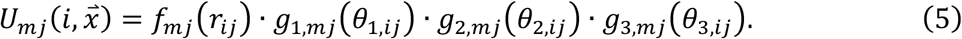

Here, *r*_*ij*_ is the distance between *B*_*i*_ and *C*_*α, j*_, and *θ*_*k,ij*_ (*k* = 1, 2, 3) are three angles involving the neighboring sugar, bases, and *C*_*α*_ particles as defined in Figure 2B. The *θ*_1,*ij*_ represents the orientation of the sugar-to-base *B*_*i*_ bond relative to the amino acid *C*_*α, j*_. The *θ*_2,*ij*_ monitors the orientation of the DNA strand relative to the amino acid *C*_*α, j*_. The *θ*_3,*ij*_ indirectly monitors the sidechain orientation of the amino acid *C*_*α, j*_ relative to the base *B*_*i*_. The modulating function of *r*_*ij*_ is defined as a Gaussian centered at the native value *r*_0,*mj*_ with the width of *σ*:

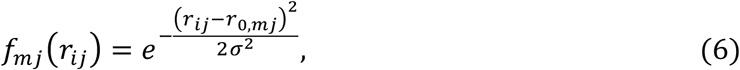

and the modulating function for *θ*_*k,ij*_ is taken from the form used in the 3SNP.2 DNA model^24^ as defined in the following:

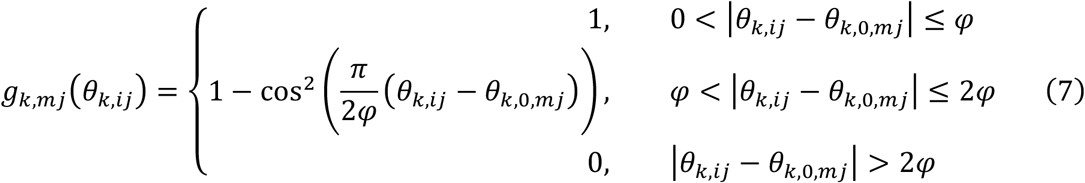

where *θ*_*k*,0,*mj*_ are the native values of *θ*_*k,mj*_ (*k* = 1,2,3) in the PDB structure, and the general parameter *φ* determines the width of the bell-shape modulating function (Supplementary Figure S1).

### Molecular Dynamics Simulations

The above-described PWMcos model was implemented in the MD package CafeMol^34^ and was applied to study protein-DNA binding by conducting Langevin dynamics simulations at temperature *T* = 300*K* (see Supporting Information for more details). Although the mapping to real time is inherently difficult, our previous estimate by comparing the diffusion coefficient suggests that one MD step roughly corresponds to an order of 1 ps.^35^

For the PWMcos function, we first run simulations to calibrate the parameters *γ* and *ε*′ in Equation (3) by matching the simulated dissociation constant (*K*_*d*_) to the experimental value. In these simulations, the distance between the center-of-mass (COM) of protein and DNA was constrained to be smaller than 100Å, which set the effective concentration. The dissociation constant (*K*_*d*_) was then estimated from the simulations by computing the populations of the bound/unbound states and compared with experimental values (more details are described in Supporting Information). For the reference structures of PU.1 and TATA-box binding protein (TBP) binding to their consensus DNAs, we used the PDB entries 1PUE^36^ and 1CDW,^37^ respectively. The PWM for both proteins were downloaded from JASPAR.^38^

After calibration, we performed simulations for *Mus musculus* PU.1 and *Homo sapiens* TBP binding to different DNA sequences including artificially designed sequences and real genome sequences. The genome sequences for PU.1 binding was downloaded from the database PAZAR.^39^ Whereas for TBP, we randomly chose 51 TATA-containing promoters, 50 TATA-less promoters, and 50 coding region sequences from the human genome.^40^ The criteria we used to classify TATA-containing or TATA-less promoters is the regular expression “TATA[at]A[at][ag]”.^41^ A full list of all the studied DNA sequences, as well as simulation time and the number of individual trajectories for each sequence, can be found in Supplementary Table S1.

All MD simulations were performed with the CafeMol package.^34^

### Analysis of Simulation Results

To quantitatively analyze the sliding behavior of protein on DNA, we recorded the indices of DNA base pairs (*I*_1_, *I*_2_, …, *I*_r_) that formed contacts with proteins in every snapshot and used the averaged DNA index, 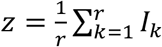, as the coordinate that represents the protein binding position on DNA. To represent the relative orientation of protein with respect to DNA, we calculated the angle between two vectors, one connecting two selected *C*_*α*_ beads, the other connecting the bases at the two ends of DNA. Specifically, we chose Lys1 and Lys78 for the orientation vector for PU.1.

To investigate the interface that proteins used to bind DNA, we used two different strategies. For PU.1, we calculated the “native-ness” of the DNA-binding protein interface. Practically, we first determined a set of amino acids that form contacts with DNA in the PDB structure, *S*_*nat*_ = {*C*_*α,l*_|*C*_*α,l*_ forms contact with DNA}, whose number of elements is denoted by 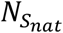. During the simulations, we then counted the number of *C*_*α*_ beads (*C*_*α*_ ∈ *S*_*nat*_) that formed contacts with DNA beads in every snapshot, denoted by *n*_*snapshot*_. The native-ness was then defined as 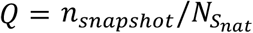.

Different from the native-ness estimation for PU.1, we employed clustering and principal component analysis for TBP-DNA binding. We first vectorized the indices of the interface residues that TBP used to bind DNA during the MD simulations (sampled every 10^6^ MD steps in 20 different 10^9^-step MD trajectories). The Euclidean distance matrix was constructed based on the chosen snapshots and was then used as the metric for the DBSCAN clustering analysis. To visualize the clustering results, we projected the vectorized snapshots onto the two-dimensional profile of the first two principal components (PC1 and PC2). Additionally, the analyzed clusters were used as the training set for the support vector classification of all the structures sampled from MD simulations. All the analysis methods mentioned here were implemented with the Python library scikit-learn.^42^

## RESULTS

### Determining Parameters in the PWMcos Functions

As described in the Methods section, we have two parameters in the distance/angle dependent modulating functions of the PWMcos (*σ* in Equation (6) and *φ* in Equation (7)). These two parameters control the shape of the modulating functions (see Supplementary Figure S1) and will consequently affect the accuracy of the energy modeling. Here we determined these parameters by considering two anticipated properties of our model. On the one hand, our model was designed for proteins to recognize DNA sequences at the resolution of 1-bp specifically; we require the sequence-specific energy to be sensitive enough to detect the binding position changes by one bp (Figure 3A). Therefore, the parameters (*σ* and *φ*) should be sufficiently small so that, when a protein is shifted by one bp along the duplex DNA, the value of the modulating function change significantly (Supplementary Figure S1A and B). On the other hand, because we represent the sequence specificity by the PWM-based factor *ε*_*m*_(*b*_*i*_) separate from the modulating function 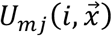 in Equation (2), we want to minimize the change in the value of the modulating functions upon a base pair mutation. Note that in the SPN.2C model the location of the base particle relative to the backbone particles changes slightly upon the mutation. Therefore, we should set the parameters (*σ* and *φ*) large enough to tolerate these changes so that the modulating function has similar value in the mutant.

**Figure 3.**
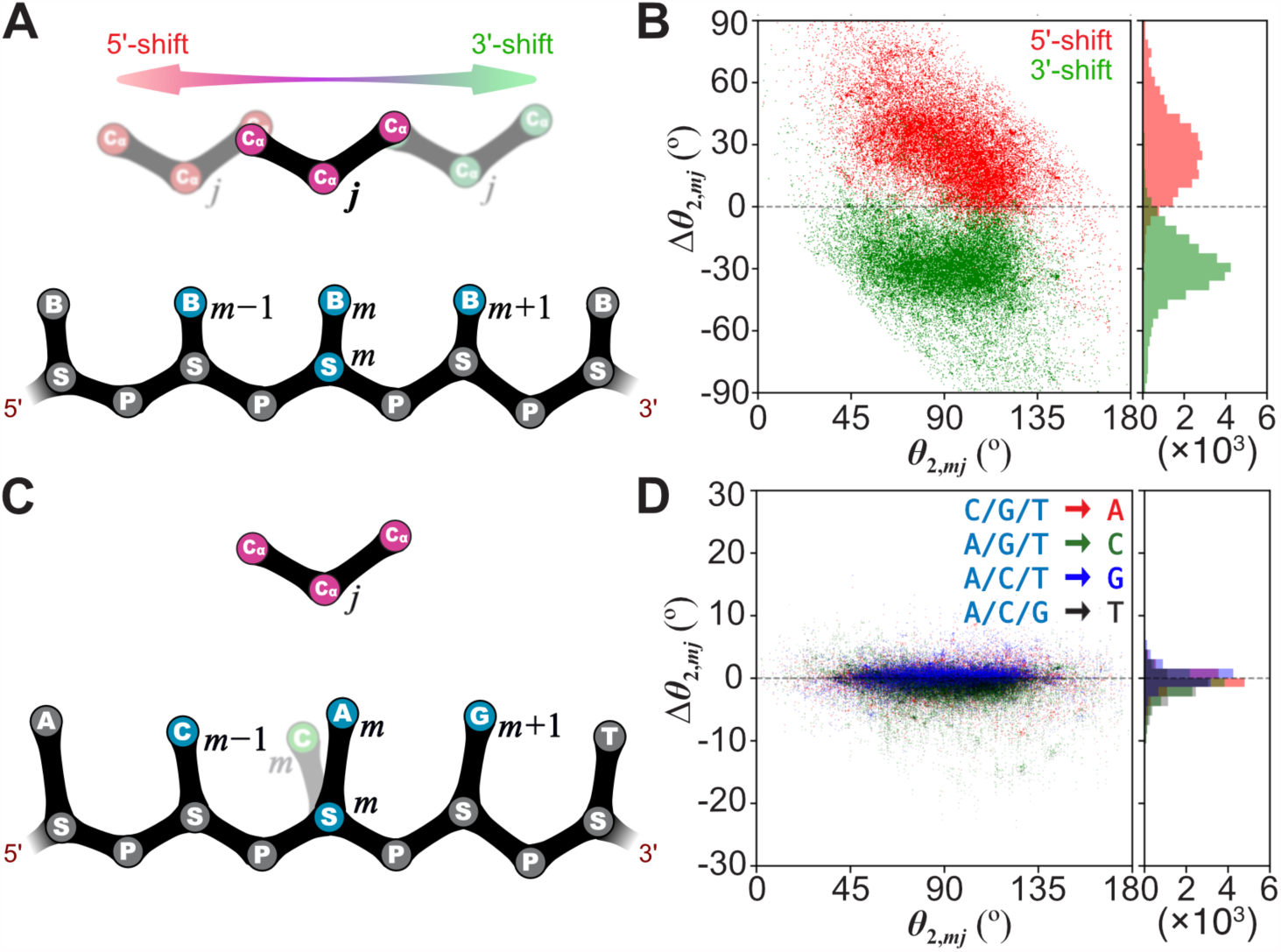
Translation and mutation tests to determine parameters in the modulating functions. (A) Schematic plot of the 1-bp shifts performed on the local structures of a protein-DNA complex. For every pair of *B*_*m*_ and *C*_*αj*_, protein residues (*C*_*αj*_ and *C*_*α*(*j*±1)_) were translocated by one base-rise (≈ 3.4Å) in both 5’ (red) and 3’ (green) directions while keeping the DNA structure fixed. (B) Scatter plot of Δ*θ*_2,*mj*_ (changes in *θ*_2,*mj*_ upon 5’-shift (red) and 3’-shift (green)) versus *θ*_2,*mj*_, calculated for the 3322 protein-DNA complex structures. The one-dimensional distributions of Δ*θ*_2,*mj*_ is shown in the right panels. (C) Schematic plot of the mutation introduced to DNA base. For every pair of *B*_*m*_ and *C*_*αj*_, *B*_*m*_ was mutated to three base types other than the native one (from A to C in the cartoon), which results in the changes of values of *r*_*mj*_ and *θ*_*k,mj*_ (*k* = 1, 2, 3). (D) is the same as (B) but for *θ*_2,*mj*_ tested with mutations illustrated in (C). The dot color represents different mutation types.

We performed statistical analysis over 3322 structures taken from the protein-DNA complex structure database NPIDB.^43^ For each pair of amino acids and bases in the complex structures, we carried out translation tests on the CG modeled *C*_*α*_ beads in both 5’-end and 3’-end directions (Figure 3A). This was executed by calculating the best-fit translational matrices from the native bases (*B*_*m*−1_, *B*_*m*_, and *B*_*m*+1_) to the 1-bp shifted ones (*B*_*m*−2_, *B*_*m*−1_, and *B*_*m*_ for the 5’-end shifts, and *B*_*m*_, *B*_*m*+1_, and *B*_*m*+2_ for the 3’-end shifts) and then applying these translations to the coordinates of the *C*_*α*_. We then calculated the changes in the distances (Δ*r*_*mj*_) and the angles (*Δθ*_*k,mj*_, *k* = 1, 2, 3) upon the translations. As an example, Figure 3B plots Δ*θ*_2,*mj*_ against *θ*_2,*mj*_ for all the tested *m*, *j* pairs. When a protein is shifted by one bp from its native binding position, *θ*_2,*mj*_ redistributes with an average deviation of 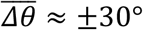. As have been discussed above, the parameter *φ* should be small enough for the modulating function *g*_2,*mj*_(*θ*_2,*mj*_) to detect the changes in *θ*_2,*mj*_. Therefore, the peak of ∆*θ*_2_ distribution roughly determines the upper limit of *φ*, namely, *φ* < 30°. The same translation tests and analyses were also carried out for *r*_*mj*_, *θ*_1,*mj*_, and *θ*_3,*mj*_, of which the results are in Supplementary Figure S1C-E.

To determine the lower limit of the parameters, we performed mutation tests for the same set of protein-DNA complex structures (Figure 3C). In a mutation test, the base that forms native contact with *C*_*α, j*_ was replaced by one of the other three base types (for instance, *b*_*m*_ = *A* → *C*, in Figure 3C). The differences in *r*_*mj*_ and *θ*_*k,mj*_, *k* = 1, 2, 3 upon the mutations were calculated and plotted against their native values (see *θ*_2,*mj*_ in Figure 3D and *r*_*mj*_, *θ*_1,*mj*_ and *θ*_3,*mj*_ in Supplementary Figure S1F-H). Figure 3D reveals that most mutations do not alter the *θ*_2,*mj*_ angle more than 5°. As we discussed above, if the parameter *φ* is large enough, *θ*_*k,mj*_ (*k* = 1, 2, 3) variations caused by mutations will result in only tiny differences in *g*_*k,mj*_(*θ*_*k,mj*_) values, which suggests *φ* > 5° in the case of *θ*_2,*mj*_. The same discussion also applies to determining the other parameters (*r*_*mj*_, *θ*_1,*mj*_ and *θ*_3,*mj*_). Combining all the results above, we found *σ* = 1.0Å and *φ* = 10° to be acceptable values satisfying both requirements in the translation and mutation tests.

### Evaluating the sequence-specificity of *E*_*PWMcos*_

Figure 4 shows a simple evaluation of the sequence-specific protein-DNA interaction energy (*E*_*PWMcos*_) for the ETS domain of protein SAP-1 as an example. The 5-bp PWM and the X-ray structure of SAP-1-DNA complex (PDB entry: 1BC8)^44^ are shown in the insets of Figure 4A. Based on the PDB structure, we introduced all possible mutations to the 5-bp DNA sequence and then calculated the sequence-specific interaction energies employing both the direct PWM estimation (*E*_*PWM*0_) and our new model (*E*_*PWMcos*_). We computed *E*_*PWM*0_ by summing up the products of the corresponding elements of the PWM and the sequence matrix:

**Figure 4.**
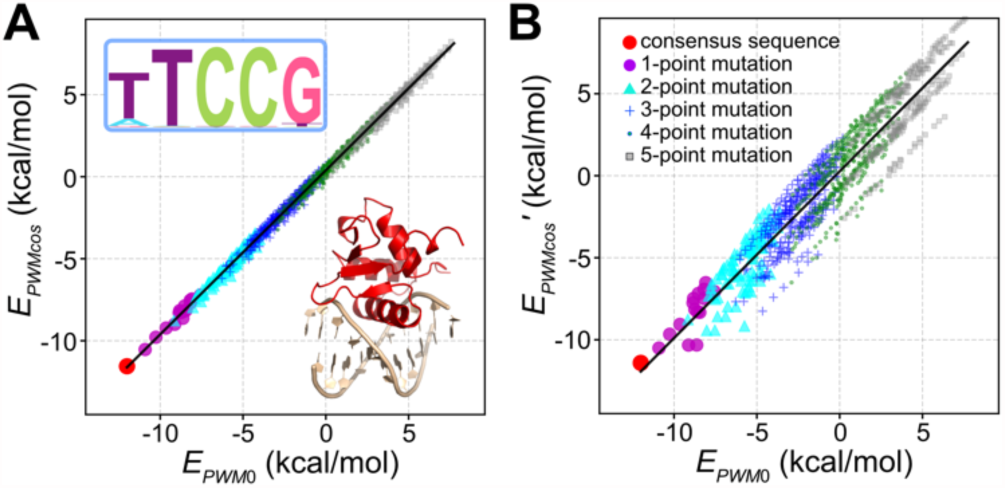
Comparison of the energy calculated by our new model (*E*_*PWMcos*_) with the PWM energy estimation (*E*_*PMW*0_) for 4^5^ DNA sequences for SAP-1 protein. We test the full angle-dependent energy function (A) as well as a simpler isotropic energy function (B) as a control. The PDB structure (PDB: 1BC8) and the 5-bp sequence logo are depicted in the inset of (A). Color and shape of dots indicate the different number of mutations from the consensus sequence.

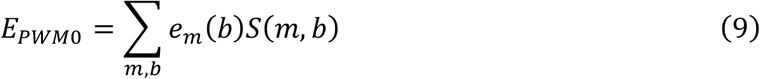

where *e*_*m*_(*b*) is the element of the PWM (defined in Equation (4)), and *S*(*m*, *b*) is the element of the sequence matrix. *S*(*m*, *b*) equals to 1 if base at position *m* has the type *b*, and 0 otherwise.

As can be seen in Figure 4A, *E*_*PWMcos*_, and *E*_*PWM*0_ for all the native and mutated DNA sequences have a good correlation (with Pearson’s correlation coefficient *r* = 0.997). This demonstrates the ability of our model to provide binding specificity estimation for a given structure as accurate as the PWM. As a control, we also computed *E*_*PWMcos*_′ for all the native and mutated structures, which was essentially the same as *E*_*PWMcos*_, except that we took off all the angle-dependent modulating functions from Equation (5): 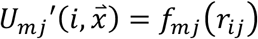. We found that without the *g*_*k,mj*_(*θ*_*k,ij*_) (*k* = 1, 2, 3) factors, *E*_*PWMcos*_′ gave poor estimations for mutated sequences (Figure 4B). Similarly, the removal of any one *g*_*k,mj*_(*θ*_*k,ij*_) (*k* = 1, 2, 3) factor resulted in significantly poorer correlation than the case of the complete function in Equation (5). These results highlight the necessity of introducing the angle-dependent modulating functions in our model, which to some extent reflects the angle-dependent hydrogen-bonding physical basis of the protein-DNA sequence-specific recognition.

We carried out similar analyses for several other proteins, which belong to a variety of different TF families. The results are shown in Supplementary Figure S2. For all the proteins we tested, our model could reproduce the relative binding affinities for protein binding to the native and mutated DNA sequences.

### PU.1 moves on rugged energy landscape and flips occasionally

Next, applying the PWMcos model, we performed CG MD simulations for two DNA-binding proteins with DNA sequences; one recognizes the major-groove of DNA (PU.1), and the other recognizes the minor-groove (TBP).

The first target, PU.1, is an ETS-family TF that plays multiple essential roles in processes such as hematopoiesis and B-cell development.^36,45–48^ For *Mus musculus* PU.1, sequence-specific DNA binding affinities^47^ and the complex structure with its consensus DNA^36^ have been characterized experimentally (Figure 5A). The central part of the consensus sequence is an 8-bp long piece, “GAGGAAGT”. From the atomistic structure of the PU.1-DNA complex, we see that PU.1 mainly uses one of its *α*-helix to recognize bases in the major groove of DNA, while a short loop forms some contacts with sugar and phosphate particles in the minor groove. Using the 8-bp long PWM and the PDB structure, we set up the PWMcos function and performed preliminary simulations for PU.1 binding to the consensus sequence. Comparing the simulated binding/dissociation rates with the experimental characterized *K*_*d*_,^47^ we calibrated the two parameters in Equation (3) to be *γ* = 2.5, *ε*′ = 0.4 kcal/mol for PU.1 (see Supporting Information for details).

**Figure 5.**
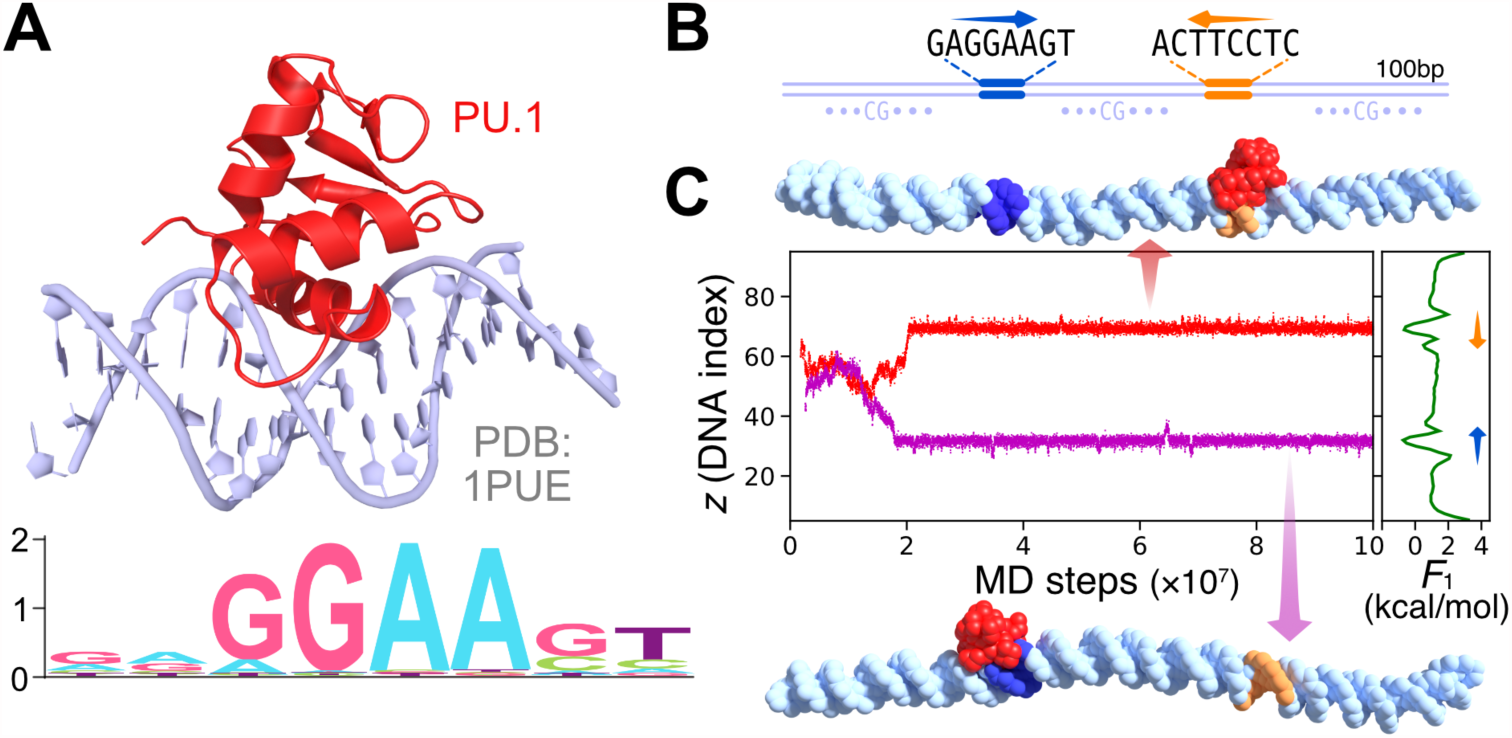
MD simulations of a transcription factor PU.1 bound to a designed 100bp DNA sequence. (A) Crystal structure of the PU.1-DNA complex (PDB entry: 1PUE) and the sequence logo of PU.1 target DNA. (B) The designed DNA sequence consists of repeating CG dinucleotide steps with two pieces of PU.1 consensus sequences (GGAAGT) inserted at indices 31-36 (blue) and 65-70 (yellow, in the opposite direction). (C) Two representative trajectories (purple and red curves) of simulated PU.1 sliding motions. Snapshots of PU.1 bound to the first (blue) and the second (yellow) consensus targets are taken from the two trajectories as indicated by the arrows. The one-dimensional free energy profile (*F*_1_) along the binding position of PU.1 on DNA (*z*) is plotted on the right panel.

With the calibrated parameters, we simulated PU.1 binding to a designed 100-bp DNA, which includes two oppositely-oriented consensus sequence elements at DNA indexes 33 and 66 in the background of poly-CG repeat sequence (Figure 5B). Figure 5C shows two representative time series of PU.1 binding position (*z*) on the designed DNA sequence. After the initial binding and searching along DNA, PU.1 was able to find the consensus target at DNA index 33 in one trajectory (purple curve). In the other trajectory (red curve), PU.1 was finally bound to the different consensus sequence at DNA index 66. As in Figure 5C, snapshots taken from these trajectories confirmed that PU.1 correctly recognized the two consensus targets using the binding pattern identical to that in the PDB structure, with the fraction of formed native contacts *Q* = 0.925 (upper structure) and *Q* = 0.950 (lower structure). Also, the one-dimensional free energy profile obtained from 20 trajectories further demonstrates that the simulated PU.1 has a roughly equal preference for the two consensus target sequences (green curves in the central right panel of Figure 5C). These results show the capability of our model to simulate the sequence-specific protein-DNA binding in the presence of more than one consensus targets.

Next, we started to explore the target search process of PU.1 on a fragment of genomic DNA sequences. In contrast to the designed homologous background of poly-CG, we expected more occurrences of pseudo binding sites in genomic sequences. We performed simulations of PU.1 binding to a 51-bp dsDNA from the *Mus musculus* genome, which was taken from the database PAZAR (Supplementary Table S1, the label “PU.1 RS0000277”). This 51-bp DNA fragment contains a recognition element at 31-38 “G_31_A_32_G_33_G_34_A_35_A_36_c_37_T_38_” (upper characters represent the consensus bases). During the simulations, we monitored the binding position of PU.1 on DNA (*z* coordinate), as well as the relative orientation of PU.1 on DNA, which was represented by the angle *ω*_PU.1-DNA_ (defined in Figure 6A). Figure 6B plots a representative time course of PU.1 (*z* coordinate), in which the orientation of PU.1 was represented by the color (black for *ω*_PU.1-DNA_ > 90° and yellow for *ω*_PU.1-DNA_ < 90°). Clearly, the trajectory reached and stayed at the recognition element at 31-38-th bp with the correct orientation (see Supporting Information Movie S1). Notably, other than this most populated recognition element, we also observed several modest-affinity sites, located around 17-24-th bp, and 42-45-th bp. By comparing the sequences of these DNA pieces to the consensus one, we found that each of these secondary binding sites has some sequence overlap with the consensus one. For example, the sequence around DNA indices 42-45 is “a_40_A_41_G_42_G_43_g_44_g_45_t_46_T_47_”, roughly half of which fits the consensus sequence. PU.1 stays at 42-45-th bp site for ~10^7^ MD steps. We also find that PU.1 bound on DNA occasionally changes its orientation without dissociating from the DNA.

**Figure 6.**
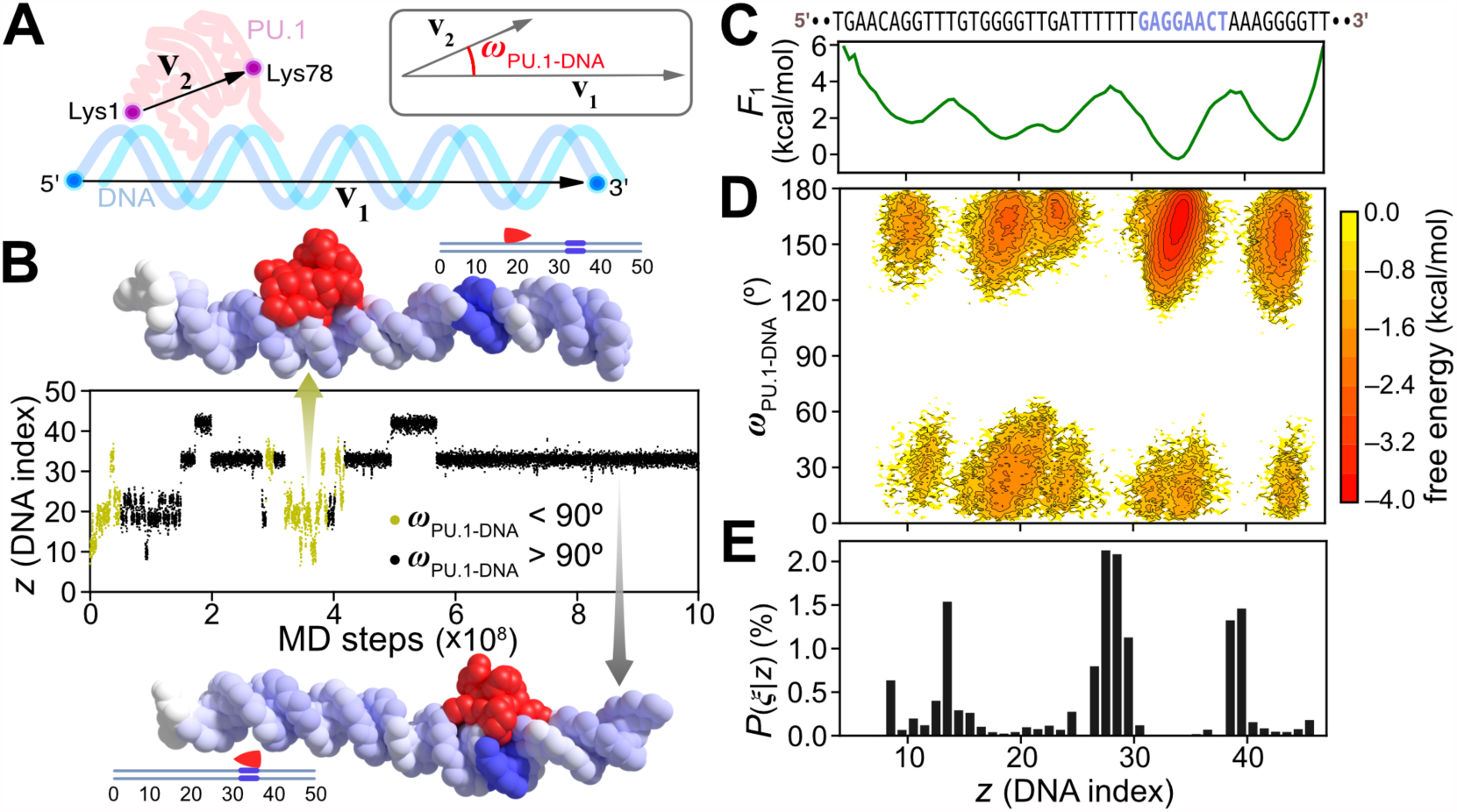
MD simulation of a transcription factor PU.1 bound to a 51-bp genomic sequence. (A) To monitor the orientation of PU.1 with respect to DNA, we employed an angle *ω*_PU.1-DNA_, defined by the two vectors, **v**_1_ and **v**_2_. (B) A representative time course of PU.1 position on DNA (*z*). The dot color represents the value of *ω*_PU.1-DNA_, yellow for acute angles and black for obtuse angles. Two snapshots showing PU.1 binding to the pseudo-target sequence (upper) and the consensus sequence (lower) are depicted. (C) Free energy profile along *z* (*F*_1_). The DNA sequence is listed at the top with the 8-bp most near-consensus sequence “G_31_A_32_G_33_G_34_A_35_A_36_c_37_T_38_” in blue. (D) Free energy surface projected onto the 2D plane of *z*-*ω*_PU.1-DNA_. (E) The probability of PU.1 to flip while binding at *z* (*P*(*ξ*|*z*)). (C-E) share the same horizontal axis (*z*).

To quantitatively compare the simulated binding specificities to the strong and weak binding sites, we computed the free energy surfaces (Figure 6C) based on data gathered from 20 independent trajectories (Supplementary Figure S3 and S4). From the free energy surface projected on the axis of PU.1 binding position (*z*), we found that PU.1 binds to the “G_31_A_32_G_33_G_34_A_35_A_36_c_37_T_38_” region with the lowest free energy, which is roughly 1.5kcal/mol stronger than the above mentioned secondary high-affinity sites. Whereas the free energy barrier near the consensus target (*z* ≈ 28) is as large as ~4kcal/mol (Figure 6C). More detailed analysis showed that although PU.1 had distinct affinities to different DNA sequences, the DNA-binding interface of PU.1 was indistinguishable on the various binding sites (Supplementary Figure S5B). These results show that our model can be used not only to simulate the protein recognition of multiple binding sites on DNA but also to distinguish the subtlest binding affinity differences between them.

We next focus on the relative orientation of PU.1 during its sliding on DNA. Based on 20 individual 10^9^-step MD trajectories, we analyzed the distribution of the angle *ω*_PU.1-DNA_ when PU.1 binds to different positions on DNA. Figure 6D shows the two-dimensional free energy surface on *z* and *ω*_PU.1-DNA_. We found that while binding to DNA, PU.1 adopted two orientations, each corresponded to *ω*_PU.1-DNA_ > 90° or *ω*_PU.1-DNA_ < 90°, respectively. The global minimum on the surface located at 32 < *z* < 35, 150° < *ω*_PU.1-DNA_ < 180°, which coincided with the conformation of PU.1 binding to the “G_31_A_32_G_33_G_34_A_35_A_36_C_37_T_38_” element using the same orientation as in the PDB structure (where *ω*_PU.1-DNA_ ≈ 159°). Yet, PU.1 also had a minor probability to accept an opposite orientation while binding to this region (Figure 6D). A more detailed analysis showed that when PU.1 was bound to a different area, the relative ratio between the two orientations also changed (Supplementary Figure S6).

In addition to the static probability distributions of PU.1 orientations, we also analyzed the transitions between the two binding orientation states. We first determined the flipping transitions from the MD trajectories by finding the adjacent frames in which PU.1 had different orientations ((*ω*_PU.1-DNA_(*t*) − 90°) × (*ω*_PU.1-DNA_(*t* + 1) − 90°) < 0). We denote the reorientation event as *ξ* and the corresponding PU.1 binding position as *z*_*ξ*_. By calculating the histogram of *z*_*ξ*_,*P*(*z*_*ξ*_), and normalizing it with the histogram of *z* from all the frames in the simulations (*P*(*z*)), we obtained the conditional probability distribution of the reorientation events (*P*(*ξ*|*z*) = *P*(*z*_*ξ*_)/P(z)). The results are shown in Figure 6E. Interestingly, we see clear peaks at DNA indices 14, 27-28, and 38-39. Combined with the free energy surfaces in Figure 6C and 6D, we found that these peaks correspond to the DNA sequences that have weaker affinities to PU.1. These results suggest that the PU.1 flip happens more frequently when the protein binds to non-consensus sequences with relatively weak interactions.

### TBP has two distinct DNA binding modes

Our second target is a minor groove binding protein, TBP, which is a crucial component of the transcription preinitiation complex.^49^ TBP is well known for its ability to recognize the “TATA-box” sequence (“TATA[at]A[at][ag]”)(the sequence logo in Figure 7A) and to bend DNA from the minor groove^37,50^ (the structure in Figure 7A, PDB entry: 1CDW), which is essential in the process of transcription initiation.^51^ TBP uses a surface mainly composed of *β*-sheet and loops to insert them into the distorted DNA minor groove. Although the TBP has a highly symmetric structure in the interface, the amino acid sequence is not symmetric, and consequently the consensus sequence of TBP is different from a palindrome (Figure 7A). It is interesting to note that the DNA-binding interface of TBP, which locates at the concave side of the protein structure, consists of a modest number of positively charged residues. Contrarily, the most positively charged region is on the convex surface of TBP (Supplementary Figure S7). We used the 8-bp PWM and the PDB structure 1CDW to construct the PWMcos interaction.

**Figure 7.**
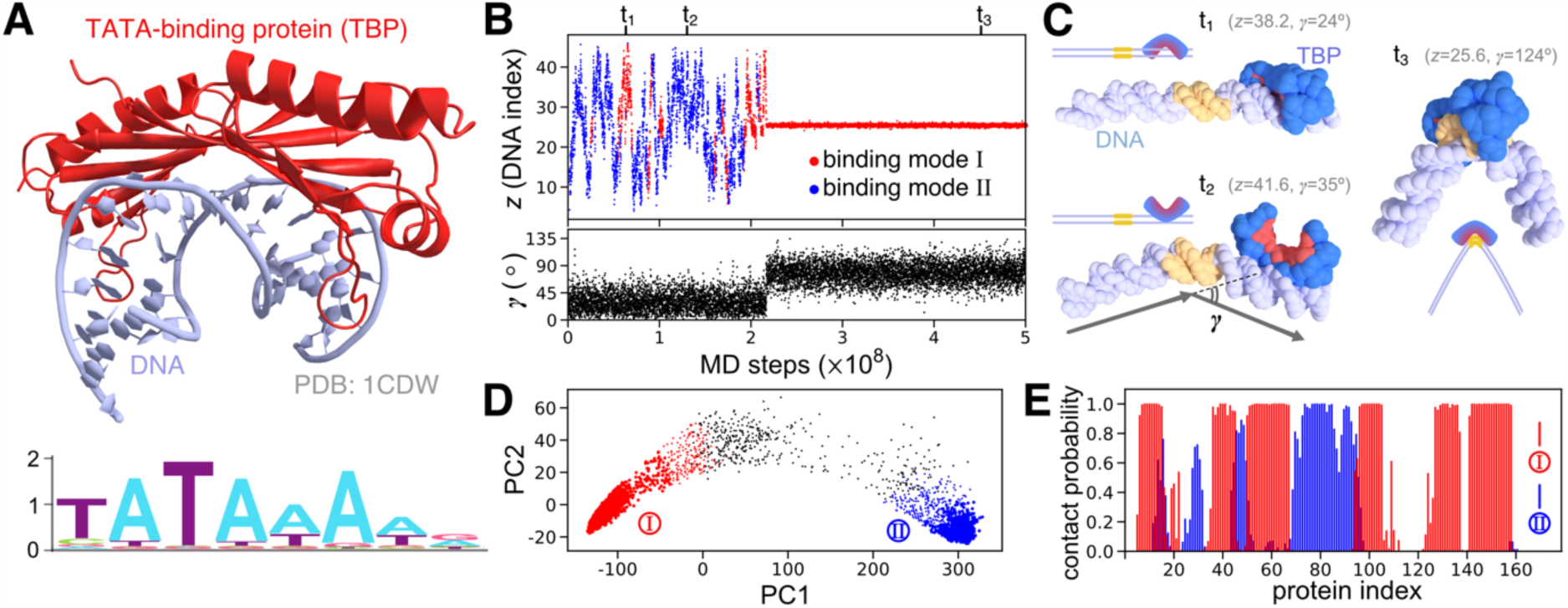
MD simulation of the TATA-box binding protein (TBP) binding to a 50-bp genomic sequence. (A) Crystal structure of TBP-DNA complex (PDB entry: 1CDW) and the sequence logo of TATA-box. (B) A representative time course of TBP binding position on DNA (*z*, upper panel) and the DNA bending angle (*γ*, lower panel). (C) Three snapshots in the trajectory given in (B) at *t*_1_ = 6434, *t*_2_ = 13056, *t*_3_ = 45577 (× 10^4^ MD steps) labelled in (B). DNA is in grey with the consensus sequence in yellow. Amino acids of TBP that form the native contacts in the PDB structure are in red, otherwise in blue. (D) DNA-bound TBP configurations plotted on the first two principal components, characterize by the interface amino acids. The two largest clusters I and II are drawn in red and blue, respectively, as well as outliers in black. The point size reflects the densities of snapshots. (E) The probability that TBP amino acids form contacts with DNA bases in the cluster I (red) and II (blue).

After calibrating the two parameters in Equation (3) for TBP, we performed 20 individual 10^9^-step MD simulations for TBP binding to a 50-bp DNA sequence taken from the human genome (Supplementary Table S1, the label “TATA-containing 0”). In each of these simulations, the initial position of TBP was randomly set around the duplex DNA. During the simulations, we tracked the sliding of TBP by monitoring its binding position on DNA (*z*). Considering the ability of TBP to induce sharp bending of its consensus DNA, we also monitored the DNA bending using the angle *γ* between the two vectors; one connecting the 5’-base and the center of geometry (COG) of the consensus sequence, and the other connecting the COG of the consensus sequence and the 3’-base (Figure 7C).

Figure 7B shows a representative time course of the position *z* (the top panel) and the DNA bending angle *γ* (the bottom panel) in an MD trajectory (also see Supporting Information Movie S2). The former suggests that, after the initial binding, TBP did not directly bind to the consensus sequence, but rapidly diffused along DNA until *t* = 2.17 × 10^8^, after which TBP was stably bound on the TATA-box (T_22_A_23_T_24_A_25_A_26_A_27_A_28_G_29_). The DNA bending angle *γ* showed a transition at the same time point: During the TBP diffusion, DNA is bent, on average, ~30°. On the other hand, after TBP recognized the consensus sequence, the DNA bending was markedly enhanced to ~80°, on average (Figure 7B). We also depict in Figure 7C three snapshots taken at different MD time steps in the same trajectory. In the structures at *t*_1_ and *t*_2_, TBP were bound to non-consensus regions, and DNA only underwent modest spontaneous bending. Whereas at *t*_3_, TBP recognized the TATA-box and inserted its β sheet into the minor groove, which resulted in a sharp bending of DNA. The *t*_3_ structure highly resembled the PDB structure of the TBP-consensus DNA complex, with an RMSD of 1.86Å.

Interestingly, before reaching the consensus sequence, we observed remarkable differences between the TBP-DNA binding mode at *t*_1_ and that at *t*_2_. In the first mode at *t*_1_, TBP interacts with DNA using the concave surface (red in Figure 7C), which is like the “biting” pattern in the consensus binding. In contrast, in the second mode at *t*_2_, TBP employs its convex surface to bind DNA backbone, totally exposing the consensus DNA-binding interface (red in Figure 7C). These results suggest the existence of at least two distinct binding modes of TBP. To quantify features of all possible binding modes, we performed clustering and principal component analysis to the DNA-binding interface of TBP (see Methods). The clustering analysis suggested two major clusters, which were easy to distinguish when the whole ensemble data were projected onto the first two principal components (Figure 7D). We labeled these two clusters as the mode I and the mode II. We then used the core elements in each cluster as a training set to classify all the snapshots in MD simulations by employing the support vector machine method; each structure in the time series of *z* was assigned as in the mode I (red) or mode II (blue) (Figure 7B and 7D). In combination with the representative structures shown in Figure 7C, we note that the mode I represents the binding pattern in which TBP used the consensus binding interface to interact with DNA, whereas the mode II corresponds to the case in which TBP used the convex surface to bind DNA. TBP uses mutually exclusive surfaces in the two binding modes; the DNA contact probabilities of every amino acid in TBP in the two modes are plotted in Figure 7E. Given that the convex surface of TBP has more positive charges than the concave surface (Supplementary Figure S7), the dominant interactions in the mode II is the electrostatic interactions to the DNA backbone phosphates. Whereas the amino acid residues responsible for the DNA base recognition were facing the convex surface (Figure 7C). Thus, TBP can only read DNA sequences in the binding mode I. Note that when TBP slid on DNA, both binding modes were used (Figure 7B), we thus were interested in finding out the roles the two binding modes played during TBP target searching process.

To gain a more comprehensive understanding of TBP sliding on genome sequences, we expanded the range of DNA sequences to both TATA-containing and TATA-less promoters, as well as gene-coding regions. As described in Methods and Supplementary Table S1, we randomly chose 50 different 50-bp DNA sequences in each category (in total 150 sequences) for molecular simulation studies. The simulation of TBP binding to each selected DNA was carried out for 1.5 × 10^8^ steps. Representative trajectories were shown in Supplementary Figure S8. We performed binding mode classifications to the sampled conformations as we did in Figure 7. In Figure 8A we show the fractions of the two binding modes of TBP during its target searching process on all three categories of DNA sequences (in the case of the TATA-containing promoters, we excluded all the snapshots at which TBP was bound on the TATA-box). As can be seen, while sliding on the TATA-containing promoter sequences (before binding to the TATA-box), TBP had a probability of 0.44 ± 0.04 to use the binding mode I. Whereas for the TATA-less promoter and the code region sequences, the frequencies of binding mode I were 0.32 ± 0.02 and 0.33 ± 0.02, respectively. To confirm that the higher probability of TBP to use the binding mode I while sliding on TATA-containing promoters is not because of TATA-box itself, we also conducted simulations of TBP binding to TATA-box-removed TATA-containing sequences (Supplementary Table S1 and Supplementary Figure S9). We found that even without TATA-boxes, TBP had a probability of 0.42 ± 0.03 to adopt the binding mode I on TATA-containing promoters. These results strongly suggest that compared with other genome regions, the TATA-containing promoters have some bias in the DNA sequence so that TBP tends to use more frequently the consensus binding surface to interact with DNA.

**Figure 8.**
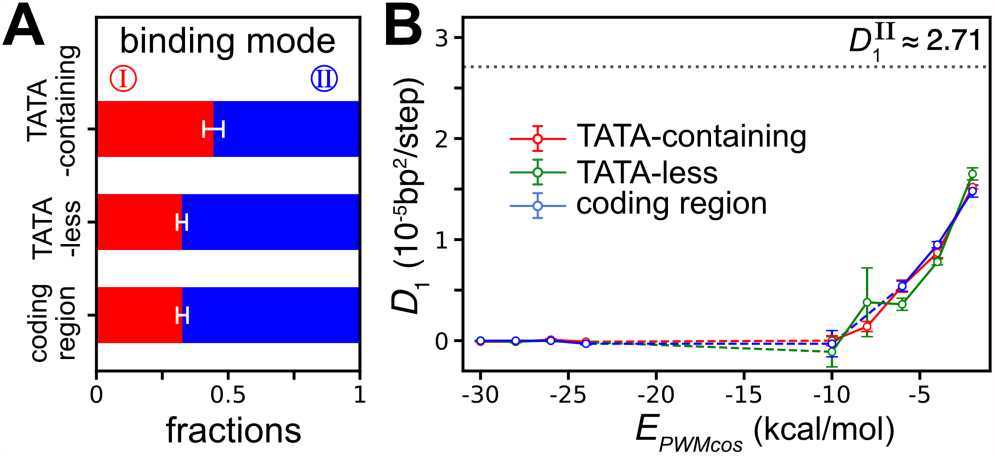
Two binding modes and diffusions of TBP on various genomic DNA sequences. (A) The fractions of the TBP binding modes in the simulations of TBP binding to 50-bp DNA sequences taken from 50 TATA-containing and 50 TATA-less promoters, and 50 gene coding regions from the human genome. For the TATA-containing promoters, we excluded the frames of TBP specifically bound to the TATA-box. (B) 1D diffusion coefficient (*D*_1_) of TBP as a function of the sequence-specific binding energy (*E*_*PWMcos*_) when TBP uses the mode I to bind the three categories of DNA sequences (red, green, and blue). The dashed lines indicate regions where *D*_1_ values are not available because of insufficient sampling. The black dotted line represents the diffusion coefficient of TBP in the binding mode II 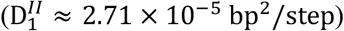.

To further investigate the sliding kinetics of TBP on DNA, we divided the MD trajectories into two groups of short pieces according to the binding mode. For each group, we analyzed the distributions of TBP sliding distances (*Δz*) during different time intervals (*Δt*) (see Supporting Information and Supplementary Figure S10), based on which we estimated the one-dimensional diffusion coefficients. We found that the diffusion of TBP in the binding mode II could be described by the simple random walk model, with the diffusion coefficient 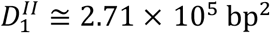/step for all the three DNA sequence categories. This result is consistent with the feature of the binding mode II that TBP binds to DNA in a sequence-nonspecific manner. In contrast, for the binding mode I, although the diffusion is apparently slower than that in the binding mode II, we found it difficult to decompose the sliding behavior of TBP into a small number of independent diffusion coefficients (see Supporting Information and Supplementary Figures S10 and S11). Instead, we tried to explore the relationship between TBP diffusion and the PWMcos interaction energy (*E*_*PWMcos*_). We performed an additional round of trajectory subdivision, based on the moving average of *E*_*PWMcos*_ (ranging from −30 to 0 kcal/mol, with a bin width of 2 kcal/mol). From the subdivided trajectories in every *E*_*PWMcos*_ interval, we computed the diffusion coefficients. In Figure 8B we show the results of computed *D*_1_ as a function of *E*_*PWMcos*_ for TBP binding to the three categories of DNA sequences. In the range of *E*_*PWMcos*_ > −10 kcal/mol, *D*_1_ changed from ~1.6 × 10^−5^ bp^2^/step to ~0 as the strength of interaction energy (*E*_*PWMcos*_) increased. When TBP was bound to DNA regions with *E*_*PWMcos*_ stronger than −10 kcal/mol, diffusion was thoroughly inhibited (*D*_1_ ≅ 0). Note that even with weak sequence-specific interactions (*E*_*PWMcos*_ ≈ −2 kcal/mol), *D*_1_ for TBP in the binding mode I was by far smaller than that in the binding mode II (Figure 8B).

In summary, these results reveal a picture of binding pattern-dependent diffusion of TBP that in the binding mode II, TBP slides in a faster, and sequence independent diffusion mode; whereas in the binding mode I, TBP adopts a slower, and energy-dependent diffusion mode.

## DISCUSSION

In the current PWMcos model, the PWM is employed to introduce the DNA sequence specificity for protein binding. However, we emphasize that our target is neither to assess the capability of the PWM modeling method nor to predict protein-DNA binding specificity but to build a model for molecular dynamics (MD) simulations to study dynamic and structural aspects of protein-DNA interactions guided by the available sequence specificity knowledge. From MD simulations, we can obtain more insights than affinity values. But, notably, the accuracy is still limited by the PWM modeling paradigm. There has been the argument about the inaccuracy of the PWM models caused by the assumption of the independence of base contributions.^52,53^ However, a recent systematic comparison of many specificity modeling algorithms^54^ showed that some newly developed variations of the simple PWM modeling^33,52,55^ had gained similar performance to more complex models.^56,57^ Note that it is easy to extend the current model to adopt different definitions of the PWM in our model; the only change to make is to substitute Equation (4) with a new form of *e*_*m*_(*b*).

How DNA-binding proteins search their target DNA sites has been extensively studied and multiple diffusion modes including 1D-sliding, hopping, intersegmental translocation, and 3D-diffusion, have been widely accepted.^15,58–60^ Here we used the PWMcos model to study the 1D-sliding process taking the DNA sequence specificity into account. For PU.1, during the sliding, its DNA-binding interface was nearly identical to that used when PU.1 binds on the consensus sequence. For TBP, we found it uses two distinct sliding modes during the search; one with the interface nearly identical to that at the consensus sequence (the mode I), and the other with an opposite side of protein surface (the mode II). In the mode I, the main contribution to binding comes from the sequence-specific interaction, whereas in the mode II, the dominant interaction is the electrostatics. These results partially explain the previous experimental observations that the nonspecific binding was affected by ionic concentration, but the consensus binding was not.^61^ Notably, the diffusion is faster in the mode II (Figure 8B). Interestingly, in genomic regions far from the consensus target, where DNA sequence bears very limited resemblance to TATA-box, TBP tends to use the diffusion mode II; while approaching the nearby areas of TATA-box target, it increases the use of the diffusion mode I (Figure 8A). This is a clear example of the two-mode model for the TF search mechanisms proposed earlier.^62,63^ By having the switchable two binding modes, TBP can diffuse more efficiently in regions far from the target site and furthermore find the target site with higher probability.

Despite the importance of TBP as a core part of the transcription initialization machinery, some basic mechanisms of TBP-DNA interactions are remaining unclear, such as the kinetics of TBP binding to the TATA-box,^64,65^ and the bending angle of nonspecific DNA upon TBP binding.^66,67^ Our simulation results suggest that while binding to sequence-nonspecific areas using the mode II, TBP does not bend DNA to the extent as in the consensus binding (Figure 7B and C), which is consistent with the demand of target search efficiency. Besides, since recent studies point out that only ~2% of human promoters contain TATA-box,^41^ the biological role that TBP plays on TATA-less promoters has been unclear. In this study we investigated the binding of TBP to TATA-containing, TATA-less promoters, and coding region DNA sequences taken from the human genome. Interestingly, TBP takes the mode I more frequently in the TATA-containing promoters relatively to the TATA-less promoters and gene-coding regions (Figure 8A). We consider that this difference is caused by the following two reasons: firstly, in the coding region and TATA-less promoters, fewer occurrence of pseudo targets results in a higher fraction of the binding mode II, in which TBP diffuses faster; secondly, the pseudo consensus binding sites in the coding region and TATA-less promoters are relatively weaker than in the TATA-containing promoters (Supplementary Figure S12), which also results in faster diffusion of TBP in these regions (Figure 8B). Indeed, several recent studies have reported differences in sequence compositions and structural properties of TATA-containing and TATA-less promoters.^41,68,69^ Here, our results provided a more dynamic view of the consequences of these sequential and structural differences.

In this study, we focused on the sequence-specific protein-DNA interaction via the DNA bases, excluding the DNA backbone contribution. In some important cases, however, proteins make significant interactions to the DNA backbone atoms in a sequence-nonspecific manner. Some notable examples are helicase interactions with the translocated DNA and histone interaction with nucleosomal DNA. In these cases, protein-DNA hydrogen bonds make these interactions highly orientation-dependent. We can easily extend our current approach to these cases. Indeed, as a separate work, we recently implemented histone-DNA hydrogen bonds in a very similar manner which enables to stabilize nucleosomal DNA at high precision.^70,71^

## CONCLUSION

In conclusion, we developed PWMcos, a new model for the protein-DNA sequence-specific interactions in coarse-grained MD simulations. The model made use of experimental results about the protein-DNA binding affinities (PWMs) and the complex structure data. With this modeling, we succeeded to reproduce the free energy surfaces of proteins binding on genome DNA sequences. Additionally, we revealed exciting mechanisms of protein diffusion kinetics, such as the sequence-dependent reorientation of PU.1 and the transition between different binding modes of TBP. We predicted that the binding modes are related to different diffusion modes, which could be tested by single-molecule experiments. The PWMcos is general and can be applied to any protein-DNA interactions only if their PWMs and complex structure data are available. The PWMcos model is publicly available in the latest version of CafeMol.

## ASSOCIATED CONTENTS

### Supporting Information

The following files are available free of charge.

More details about MD simulations and discussions about diffusion mode analysis of TBP, as well as all the Supplementary Figures. (PDF)

Supplementary Movies S1, S2. (MPG)

### Funding Sources

This work was supported by JSPS KAKENHI [25251019 to S.T., 16KT0054 to S.T.,16H01303 to S.T.]; MEXT as ‘Priority Issue on Post-K computer’ (to S.T.); RIKEN Pioneering Project ‘Dynamical Structural Biology’ (to S.T.). The funders had no role in study design, data collection and analysis, decision to publish or preparation of the manuscript. Funding for open access charge: MEXT ‘Priority Issue on Post-K computer’.

### Conflict of Interest

The authors declare no conflicts of interest.

## ACKNOWLEDGEMENT

We thank Giovanni B. Brandani for many helpful discussions.

---

**For Table of Contents use only**

Dynamic and Structural Modeling of the Specificity in Protein-DNA Interactions Guided by Binding Assay and Structure Data

By Cheng Tan and Shoji Takada

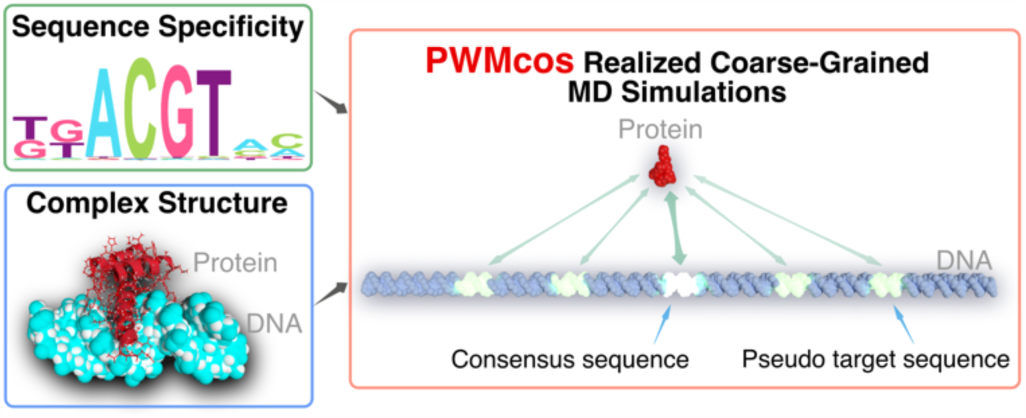

