## Supplementary Materials for "Dynamic and Structural Modeling of the Specificity in Protein-DNA Interactions Guided by Binding Assay and Structure Data"

March 23, 2018

### 1 Supplementary Methods

#### 1.1 Details of Protein AICG2+ model

The coarse-grained (CG) model we used for protein was the AICG2+ model[1], which was developed on top of the original Gō model[2], with enhancements based on matching the fluctuations in CG simulations to the atomic results produced with the AMBER force field[3]. The AICG2+ energy potential was defined as the following[1]:

$$\begin{aligned} V_{AICG2+}(R|R_0) &= V_{local} + V_{G\bar{o}} + V_{exv} \\ &= \sum_i k_{b,i}\big(r_{i,i+1} - r_{i,i+1,0}\big)^2 + V_{local}^{flp}(R) \\ &+ \sum_i \varepsilon_{loc2,i} \exp\Big(-\frac{\big(r_{i-1,i+1} - r_{i-1,i+1,0}\big)^2}{2W_{i-1,i+1}^2}\Big) + \sum_i \varepsilon_{loc3,i} \exp\Big(-\frac{\big(\phi_i - \phi_{i,0}\big)^2}{2W_{\phi,i}^2}\Big) \\ &+ \sum_{i<j-3}^{\text{native}} \varepsilon_{G\bar{o},i,j}\Bigg(5\Big(\frac{r_{i,j,0}}{r_{i,j}}\Big)^{12} - 6\Big(\frac{r_{i,j,0}}{r_{i,j}}\Big)^{10}\Bigg) \\ &+ \sum_{i<j-3}^{\text{non-native}} \varepsilon_{exv}\Bigg(\Big(\frac{\sigma_{i,j}}{r_{i,j}}\Big)^{12} - \Big(\frac{\sigma_{i,j}}{r_{i,j,cutoff}}\Big)^{10}\Bigg). \end{aligned} \tag{S1}$$

The first term in Eq.(S1) is for bonds between neighboring $C_\alpha$ particles. The third and the fourth terms are the angle and dihedral-angle potentials, respectively. In these two parts, parameter $W$s control the Gaussian widths. The fifth and the sixth terms are the classical Gō potentials for native contacts and the excluded volume interactions for non-native contacts, respectively. The second part in Eq.(S1) is a flexible local potential:

$$V_{local}^{flp}(R) = -k_B T \sum_i \ln \frac{P_\theta(\theta_i|i)}{\sin\theta_i} - k_B T \sum_i \ln P_\varphi(\varphi_i|i), \tag{S2}$$

where $P_\theta(\theta_i|i)$ and $P_\varphi(\varphi_i|i)$ are residue type dependent probability distributions of the bond angle ($\theta$) and the dihedral-angle ($\varphi$), respectively. $k_B$ is the Boltzmann constant, while $T$ is the simulated temperature. As for the excluded volume term, we employed a set of residue type dependent particle radii, as described in our previous work[4].

#### 1.2 Electrostatic Interactions

We modeled the electrostatic interactions between charged CG particles by the Debye-Hückel theory. The potential is given by:

$$V_{ele} = \frac{1}{4\pi\epsilon(T,C)} \frac{q_i q_j e^{-r_{i,j}/\lambda_D}}{r_{i,j}}, \tag{S3}$$

where $i$ and $j$ are the indices of the two charged particles, $r_{i,j}$ is the distance between them, $\lambda_D$ is the Debye length, and the dielectric constant $\epsilon(T,C)$ is a function of temperature ($T$) and solvent ionic strength ($C$)[5].

#### 1.3 Langevin Dynamics

We performed Langevin dynamics with CafeMol[6]. The equation of motion is as the following:

$$m_i \frac{d^2 r_i}{dt^2} = -\frac{\partial V_{total}}{\partial r_i} + m_i \gamma_i \frac{dr_i}{dt} + m_i \xi_i,$$

where the random noise $\xi_i$ meets:

$$\langle \xi_i(t) \rangle = 0, \quad \langle \xi_i(t) \xi_j(t') \rangle = \frac{2\gamma_i k_B T}{m_i} \delta_{t,t'} \delta_{i,j}.$$

In the above equations, $m_i$ represents the mass, $\gamma_i$ is the friction coefficient, $k_B$ is the Boltzmann constant, and $T$ is the temperature set to $300K$. For particle masses and the friction coefficent, we used the default settings in CafeMol.

#### 1.4 Calibration of Parameters for the Sequence-Specific Interactions

As we mentioned in the main text, there are two parameters, $\varepsilon'$ and $\gamma$ in the PWMcos energy function (see Eq.(3)). Based on the assumption of PWM modeling that contributions of each basepair are independent and additive, changes in $\varepsilon'$ will not affect the relative binding affinity of the protein to different DNA sequences. The reason to introduce the scale factor, $\gamma$, is that the "energy modeling" of the PWM elements we used in the current work (see Eq.(4) in the main text) in fact represents "free energy" of protein-DNA binding. As an offset against the inclusion of entropy into energy modeling, we used the factor $\gamma$ to tune the relative contributions of each base pair to the binding affinity.

In principle, for every protein of interest, we can calibrate these two parameters by running simulations and match computed thermodynamic state functions with available experimental results. To determine two parameters, we need at least two physical quantities for calibration. A reasonable combination could be the binding affinity (or equivalently the dissociation constant) for the consensus binding and the PWM. Practically, we first chose a pair of different DNA sequences (say, sequence A and B) and computed the binding free energy difference between them using PWM ($\Delta G_{PWM}(A, B) = G_{PWM}(B) - G_{PWM}(A)$). On the other hand, from MD simulations (the system should include both sequences A and B) we can compute $\Delta G_{MD}(A, B) = -k_B T \ln \frac{P(B)}{P(A)}$, where $P(A)$ and $P(B)$ are the probabilities of observing protein binding to sequence A or B in simulations, respectively. We first fixed the value of $\varepsilon' = 0$kcal/mol and determined $\gamma$ by choosing a proper value that gave $|\Delta G_{MD}(A, B) - \Delta G_{PWM}(A, B)| \simeq 0$. After this, we fixed $\gamma$ and determined $\varepsilon'$ by matching the simulated $K_D$ to the experimental values. As described above, $\varepsilon'$ controls the strength of protein-DNA interactions without affecting the sequence specificity. From simulations, we calculated the dissociation constant by the formula $K_D^{MD} = \frac{(1 - P_b)^2}{P_b V}$, where $P_b$ was the probability of the bound state, and $V$ was the volume of the simulation box. An ideal value of $\varepsilon'$ would make $K_D^{MD}$ a good approximation to the experimental value $K_D^{ex}$. Note that in some cases the ionic concentrations used in the $K_D^{ex}$ measurement experiments are too low to model with the Debye-Hückel theory in simulations. For these cases, before tunning $\varepsilon'$, we first determined a proper ionic strength in simulations by comparing the sequence-nonspecific $K_D^{MD}$ with experimental values (if the sequence-nonspecific dissociation constant is available).

In the current work, we employed experimental results of $K_D^{ex} = 453.5nM$ for PU.1 sequence-specific binding and apparent $K_D^{ex} = 2000nM$ for PU.1 sequence-nonspecific binding[7]. As for TBP, we used experimental values of $K_D^{ex} = 0.46nM$ for the sequence-specific binding and $K_D^{ex} = 340nM$ for the sequence-nonspecific binding[8]. Note that in the case of the sequence-nonspecific binding, if the experimental $K_D^{ex}$ value is provided for a single binding site, we have to translate it into apparent $K_D^{ex}$ by considering the number of possible binding sites in a simulated DNA sequence. By testing different combinations of $\gamma$ and $\varepsilon'$ values, we found that $\gamma = 2.5, \varepsilon' = 0.4\text{kcal/mol}$ worked well for PU.1 and $\gamma = 3.9, \varepsilon' = 0.0\text{kcal/mol}$ reproduced experimental behaviors for TBP.

##### 1.5 Analyzing Diffusion Modes of TBP

We performed 50 individual MD simulations for TBP binding to TATA-containing promoters, 50 for TATA-less promoters, and 50 for coding region DNA sequences (150 simulations in total). For every MD trajectory, we first divided the trajectories into shorter trajectory pieces based on TBP binding pattern (same as we showed in Fig. 7B). We then classified all the trajectory pieces into two groups, each of which corresponded to the binding pattern I or II. For these two trajectory groups we calculated the distributions of TBP sliding distance ($\Delta z$) in different time intervals ($\Delta t = 5, 10, 20, 30, 40, 50, 60, 70, 80, 90, 100 \times 10^4$ steps). The results are shown in Fig. S10.

For binding pattern II (with TBP using the nonspecific binding interface), we found that single Gaussian fitted well with the $\Delta z$ distributions, which suggested the TBP diffusion in binding pattern II was the simple random walk. We computed the one-dimensional diffusion coefficient by

$$D_1 = \frac{\text{MSD}_{\Delta t}}{2\Delta t},$$

where $\text{MSD}_{\Delta t} = \langle \big(z(t{+}\Delta t){-}z(t)\big)^2 \rangle_t$ is the mean square deviation of TBP binding position ($z$). The calculated $D_1$ values were $2.71 \pm 0.12 \times 10^{-5}\text{bp}^2/\text{step}$ for TATA-containing promoters, $2.71 \pm 0.15 \times 10^{-5}\text{bp}^2/\text{step}$ for TATA-less promoters, and $2.72 \pm 0.14 \times 10^{-5}\text{bp}^2/\text{step}$ for coding region sequences, respectively (see blue dots in Fig. S11).

For the binding pattern I (with TBP using the specific binding interface), we tried to fit the $\Delta z$ distributions with double Gaussians (single Gaussian fit is manifestly improper, see Fig. S10). For each of the three DNA sequence categories, we decomposed the sliding behavior of TBP into two modes, one with almost zero diffusion coefficient ($D_1 \simeq 0$, see the red triangles in Fig. S11), the other with finite diffusion coefficient ($D_1 > 0$, see the red circles in Fig. S11). However, $D_1$ of the fast diffusion mode of TBP on the three DNA categories was inconsistent (Fig. S11). These results showed that diffusion of TBP in the specific binding pattern could not be explained by a simple two-mode model. One possible reason could be that $D_1$ is a continuous function of $E_{PWMcos}$, and the distribution of $E_{PWMcos}$ then determines the sliding behavior of TBP on different categories of DNA sequence. Indeed, we found that different DNA sequences resulted in significantly distinct distributions of $E_{PWMcos}$ in the MD simulations (Fig. S12). Based on the assumption that $D_1$ is dependent on $E_{PWMcos}$, we re-performed the division of trajectories according to the moving average of $E_{PWMcos}$ during simulations and calculated $D_1$ in different $E_{PWMcos}$ domains. These results are described and discussed in the main text.

#### 2 Supplementary Tables

Table S1: DNA sequences used in the MD simulations. For each sequence, we list the number of individual trajectories and simulation time in each trajectory in the third column, in the format of $t \times n$, where $t$ is the number of MD steps in one trajectory and $n$ is the number of individual simulations.

| Label | DNA Sequence | Simulation Time |
|---|---|---|
| PU.1 $K_D$ nonspecific | 5'-ACGTCAGTTCTCTTTTACGT-3' | $(1 \times 10^8) \times 80$ |
| PU.1 $K_D$ specific | 5'-ACGTCACTTCCCCTTTACGT-3' | $(1 \times 10^8) \times 120$ |
| PU.1 designed | 5'-GCGCGCGCGCGCGCGCGCGCGCGAAAGGGGAAGTGCGCGCGCGCGC<br>GCGCGCGCGCGCGCACTTCCCCTTTGCGCGCGCGCGCGCGCGCGCG-3' | $(1 \times 10^8) \times 20$ |
| PU.1 RS0000277 | 5'-CTTTGAACAGGTTTGTGGGGTTGATTTTTTGAGGAACTAAAGGGGTTTTTT-3' | $(1 \times 10^9) \times 20$ |
| TBP $K_D$ nonspecific | 5'-GCGCGCGC-3' | $(1 \times 10^8) \times 180$ |
| TBP $K_D$ specific | 5'-GGAATTCGGGCTATAAAAGGGGGATCCG-3' | $(1 \times 10^8) \times 20$ |
| TATA-containing 0 | 5'-GGGGCGACGCGGCCTGCCGGGTATAAAAGCTGGGCCGGCGCGGGCCGGGC-3' | $(1 \times 10^9) \times 20$ |
| TATA-containing 1 | 5'-GAGCATATTCTTCTATAAAAATATTCTTAATTTTATTATTTTTGCCTTAT-3' | $(1.5 \times 10^8) \times 1$ |
| TATA-containing 2 | 5'-AGCAATTAGCCAGGGAATGTATAAAAGGCGTCAGGGAGACTCACTGGGCT-3' | $(1.5 \times 10^8) \times 1$ |
| TATA-containing 3 | 5'-TGATTGGCTACTTTGTTCGCATTATAAAAGGCACGCGCGGGCGCGAGGCC-3' | $(1.5 \times 10^8) \times 1$ |
| TATA-containing 4 | 5'-TGGCGGGAGCCTATAAAAGCTGGTGCCGGCGCGACCCGCGGACACACAGT-3' | $(1.5 \times 10^8) \times 1$ |
| TATA-containing 5 | 5'-AGAGCCTTGGTTAAAACCCCTATAAAAGGTTCTCATCACACTGACCTGGT-3' | $(1.5 \times 10^8) \times 1$ |
| TATA-containing 6 | 5'-CAAGACCGGTATAGCTACGTTATTATAAAAACAATGAGAATAGGGGAGTG-3' | $(1.5 \times 10^8) \times 1$ |
| TATA-containing 7 | 5'-AGCCACGCCCAGCCGAGGGACGTATAAAAGGCAGGTCTAGCAGACTAACC-3' | $(1.5 \times 10^8) \times 1$ |
| TATA-containing 8 | 5'-CTAGGGAGGATGTGGCGGGTATAAAAGCCCCACCCAGGCCAGCCGGCTCT-3' | $(1.5 \times 10^8) \times 1$ |
| TATA-containing 9 | 5'-GCGGGCGGGGCCGAACGTGGTATAAAAGGGGCGGGAGGCCAGGCTCGTGC-3' | $(1.5 \times 10^8) \times 1$ |
| TATA-containing 10 | 5'-TCTGACATGTATATAAATTCTGAGCTCTCCAAAGCCCACTGCCAGTTCTC-3' | $(1.5 \times 10^8) \times 1$ |
| TATA-containing 11 | 5'-CAGGGTATAAAAGAGGATCTGGGGCAAGGAAACTTCCAAACCTTCCAAAC-3' | $(1.5 \times 10^8) \times 1$ |
| TATA-containing 12 | 5'-GAACACCCTCGAAGTGTGTATATAAAGGCCAAAAAGGCAGCCTTGAATAC-3' | $(1.5 \times 10^8) \times 1$ |
| TATA-containing 13 | 5'-ATATATAAGAAGGAAAGGAGGAAGATGGCTTTGGGAGGAGGAGAGGGAAG-3' | $(1.5 \times 10^8) \times 1$ |
| TATA-containing 14 | 5'-TTCCTTTTAGTAGGTTCCTCATTTTACCATTTCAAACATCTTATAAATGT-3' | $(1.5 \times 10^8) \times 1$ |
| TATA-containing 15 | 5'-CTATAAAAGACATATTCAGCTCCAGCACCTCATCTGCTCTGACTTCCCCA-3' | $(1.5 \times 10^8) \times 1$ |
| TATA-containing 16 | 5'-TGGAACAGATATAAAAGGACTGTATGAGGCAAAAGAAGTCTTCTGGGAAT-3' | $(1.5 \times 10^8) \times 1$ |
| TATA-containing 17 | 5'-GCTAGTATATAAGGACTCGACCTGGGCGCCTCCAGGGAAACGCTAAGGGG-3' | $(1.5 \times 10^8) \times 1$ |
| TATA-containing 18 | 5'-AATATTTGCTATAAATAGGGCCATCCCAGGCTGCTGTCAGAATATAACAG-3' | $(1.5 \times 10^8) \times 1$ |
| TATA-containing 19 | 5'-TTCCTGTTGCCGGGACGCTATATATAACGTGATGAGCGCACGGGCTGCGG-3' | $(1.5 \times 10^8) \times 1$ |
| TATA-containing 20 | 5'-TAAAGATACATATGTATGCATTTGTGTGTGTATGTATGTGTATATAAATA-3' | $(1.5 \times 10^8) \times 1$ |
| TATA-containing 21 | 5'-CTGTGGTGAGAATTAATCTATGATGTGTTAGACTCTCCAGCTATATATAT-3' | $(1.5 \times 10^8) \times 1$ |
| TATA-containing 22 | 5'-CTATGTTATTTTAATTTTTACTATTATATATGTGTGTGTGTGTGTGTGTA-3' | $(1.5 \times 10^8) \times 1$ |
| TATA-containing 23 | 5'-CTTTAAAACAGAAGAAAAATAACTACCTTAAATTTAAAATTGTATATAAA-3' | $(1.5 \times 10^8) \times 1$ |
| TATA-containing 24 | 5'-AGGAGCCACATCTATATATATTTCTGTGGCCTATGAAAAACAGAGTGCTT-3' | $(1.5 \times 10^8) \times 1$ |
| TATA-containing 25 | 5'-CTCTTTGGACGGGATGTACTATAAATTATATATATGCTATGTCAAGCAGT-3' | $(1.5 \times 10^8) \times 1$ |
| TATA-containing 26 | 5'-AAAAAACTTCCCTGGGGGCTATAAATATCTCTCCCTGGCTAAAGATAGTA-3' | $(1.5 \times 10^8) \times 1$ |
| TATA-containing 27 | 5'-TGGGCGGGGTTATATAAGCCAGCGGCGGGCGGAAGGCGGGGCGTGGGGGT-3' | $(1.5 \times 10^8) \times 1$ |
| TATA-containing 28 | 5'-CAGTTATAAAAAGAATGCCACACTGTTCCTCAGCCTTTCCTAGGGGTGTG-3' | $(1.5 \times 10^8) \times 1$ |
| TATA-containing 29 | 5'-GGGAGGGTATATAAGCCGAGTAGGCGACGGTGAGGTCGACGCCGGCCAAG-3' | $(1.5 \times 10^8) \times 1$ |
| TATA-containing 30 | 5'-AGTTTATAAATGAAAACCAACATAACATTTCTAGCAAATAATGGGAAGAA-3' | $(1.5 \times 10^8) \times 1$ |
| TATA-containing 31 | 5'-CGCCCGAGGCTATAAAAGCATCGCCACCTGCTGCCACTAGCCAAGCCGCG-3' | $(1.5 \times 10^8) \times 1$ |
| TATA-containing 32 | 5'-AGTTCCTGGGAGATACTGTATATATGGACGGCCAAAGCATCTTCTGAGAC-3' | $(1.5 \times 10^8) \times 1$ |
| TATA-containing 33 | 5'-AAGGGGTGTATAACTGAGACTATAAAAACTCAGAGAGAAAACTCACCACC-3' | $(1.5 \times 10^8) \times 1$ |
| TATA-containing 34 | 5'-GGCCCGGGACCAGCGCCGCGGCTATAAATGGGCTGCGGCGAGGCCGGCAG-3' | $(1.5 \times 10^8) \times 1$ |
| TATA-containing 35 | 5'-ATAAAGAACCACCATATGCTAACTAACCCTCCTGATTTTATCTATATATA-3' | $(1.5 \times 10^8) \times 1$ |
| TATA-containing 36 | 5'-ATGGTCTGATATGCCCTATAAAATGTATAAAAGACATATAAGGCACCCTA-3' | $(1.5 \times 10^8) \times 1$ |
| TATA-containing 37 | 5'-AGCATATCATCAAGAAATAACTATAAAAATAGGCAACCAGCAGCTCCCGG-3' | $(1.5 \times 10^8) \times 1$ |
| TATA-containing 38 | 5'-CGGGCGGGGCCGAACGTGGTATAAAAGGGGCGGGAGGCCAGGCTCGTGCC-3' | $(1.5 \times 10^8) \times 1$ |
| TATA-containing 39 | 5'-ACATGCCACAAAGGCACAGTATAAAAACGGTGGGAATCAGAGCACTTCAG-3' | $(1.5 \times 10^8) \times 1$ |
| TATA-containing 40 | 5'-AAAAGGCCCTATATAAGTTCGGAGTATTTATAATAATTATGATTATTATT-3' | $(1.5 \times 10^8) \times 1$ |
| TATA-containing 41 | 5'-GTCTGTCTGTCTTATAAATAATCAGAGTTTCCAAATGAACTGTAGTTTGT-3' | $(1.5 \times 10^8) \times 1$ |
| TATA-containing 42 | 5'-AGCTTATAAAAGCTCTTCATTTTATAACTTTCAGGAGGATCTCATGTATC-3' | $(1.5 \times 10^8) \times 1$ |
| TATA-containing 43 | 5'-ACGTCACCCCGGGCTTTATAAAAGGGATGTGATCAGGGAGCTGGGGAGAA-3' | $(1.5 \times 10^8) \times 1$ |
| TATA-containing 44 | 5'-AGCCTGACCAACATGGTGAAACCCCGTCTCTACTAAAAATATAAAAATTA-3' | $(1.5 \times 10^8) \times 1$ |
| TATA-containing 45 | 5'-CTAAGCCTCATCCATCACTGATCAGTCTTCACGTATAAAAGTAAGGATTT-3' | $(1.5 \times 10^8) \times 1$ |
| TATA-containing 46 | 5'-GCAACCTATATAAGGCTCACAGTCTGCGCTCCTGGTACACGCGCTTCAAC-3' | $(1.5 \times 10^8) \times 1$ |
| TATA-containing 47 | 5'-CCCTGAAGCCTATAAAAGCGCAGGTCCCCGCCTCGGCAGCCACGGGACAC-3' | $(1.5 \times 10^8) \times 1$ |
| TATA-containing 48 | 5'-TGTGTGTGTATATATATATATTTTGAGATGAAATCTCACTCTTTTGCCCA-3' | $(1.5 \times 10^8) \times 1$ |
| TATA-containing 49 | 5'-GGGTGTCTCAATATAAATACATCCTTCATGGTGGCCACTAGTAACGCAGG-3' | $(1.5 \times 10^8) \times 1$ |
| TATA-containing 50 | 5'-ACGTCGAGCTTCAAAAATATATATATTTTTGCAATTTTTAGGTTTTATAT-3' | $(1.5 \times 10^8) \times 1$ |

Table S1 continued...

| | | |
|---|---|---|
| TATA-less 1 | 5'-CCCCCATTGGCCAACCGGCGAGTCCCGCGCGCCACCTCCTGGGAGATGTA-3' | $(1.5 \times 10^8) \times 1$ |
| TATA-less 2 | 5'-TTTTCCAGCCGGAGTCAGGGTCAGAGGCAGCCTTTCCTACTGCGCAGGCT-3' | $(1.5 \times 10^8) \times 1$ |
| TATA-less 3 | 5'-CTTCCCGCGCATCTCTACGTACGCCCCACCCCGGTGCGCATGCGCCCGGA-3' | $(1.5 \times 10^8) \times 1$ |
| TATA-less 4 | 5'-TCCGATGCCAGGTTTCCTTAGCTTCCTTAGTGTGTCTTTGCGCTCACCAG-3' | $(1.5 \times 10^8) \times 1$ |
| TATA-less 5 | 5'-CCTCATAAAGCCCGTAACACTGGCCTGGGCTGGGGAAGCCTAGGTAAGCT-3' | $(1.5 \times 10^8) \times 1$ |
| TATA-less 6 | 5'-CAGTTGCTTCCGAACCTGACTTGGCTGGACCTCCGGTACAATAGAATTAA-3' | $(1.5 \times 10^8) \times 1$ |
| TATA-less 7 | 5'-GGGCTTGGAAACAAATGACAGGGGAAGTCCAAGAGAGGGAACTCGAACCT-3' | $(1.5 \times 10^8) \times 1$ |
| TATA-less 8 | 5'-GCGGATCCGCCTGCCCACCGCGCGTGCCCCGCCTCTCTCCCAAGAGCTAC-3' | $(1.5 \times 10^8) \times 1$ |
| TATA-less 9 | 5'-ATCAAACTGTTTGTGGTACTTCTAAACTTCGTGGAAGAGCCCTGTTTCAT-3' | $(1.5 \times 10^8) \times 1$ |
| TATA-less 10 | 5'-TGTCGTAGGGCAACCACGTAGTACTCTCTGCGCATGTGCAAAGCGCTGTC-3' | $(1.5 \times 10^8) \times 1$ |
| TATA-less 11 | 5'-CAACACCGCCTAGACCGACCGGATACACGGGTAGGGCTTCCGCTTTACCC-3' | $(1.5 \times 10^8) \times 1$ |
| TATA-less 12 | 5'-CCACAATAAAGAACCCCTGATATTTGACGCTGTGCACAAACGACTCTTTT-3' | $(1.5 \times 10^8) \times 1$ |
| TATA-less 13 | 5'-GGCCCGCGAGCTGAAGGGTGAGCTCTACTGCCTGCCCTGCCATGACAAGA-3' | $(1.5 \times 10^8) \times 1$ |
| TATA-less 14 | 5'-TGGGTTTCTTGTTACGTATGCTTGTCGAGATAGCAGAAAAAGAATGCAGG-3' | $(1.5 \times 10^8) \times 1$ |
| TATA-less 15 | 5'-CGCCCCCTCCCCGCGCGCGGAAGCCACGCCCCCGGCCGCGCGCTCGCGCT-3' | $(1.5 \times 10^8) \times 1$ |
| TATA-less 16 | 5'-AAGACCCTGGCCTCCATCTTGAAAACGGAAAATAAAAACAGCAGACCTAT-3' | $(1.5 \times 10^8) \times 1$ |
| TATA-less 17 | 5'-TCAAACTCAGTGCACTTGTTGAGCTCGTGAAACACAAGCCCAAGGCAACA-3' | $(1.5 \times 10^8) \times 1$ |
| TATA-less 18 | 5'-TACACTAGCCTTACAACGGAATTTCTTCTTGTGGAATAAAAACTGATAAT-3' | $(1.5 \times 10^8) \times 1$ |
| TATA-less 19 | 5'-CGCCGACGCTGCCCAAACAGGAGCCTGCGCATGCGCTCTGCCCTGGCAGC-3' | $(1.5 \times 10^8) \times 1$ |
| TATA-less 20 | 5'-GCTGGCTGCGGTGGCGCGCGGGCGCGGCACCCGGAAGTCGGCGGCGGTGG-3' | $(1.5 \times 10^8) \times 1$ |
| TATA-less 21 | 5'-AAAGGAAGAGTGAATAAGAGGGAGAAAATAGTTACATGTACTGTGAGTAG-3' | $(1.5 \times 10^8) \times 1$ |
| TATA-less 22 | 5'-CTTCTCCTGCCAGTGCAGGGAGCAGTGATCTTATCAGCTCAGATAGTCAT-3' | $(1.5 \times 10^8) \times 1$ |
| TATA-less 23 | 5'-AGTGGCGCGATCTCGGCTCACTGCAAGCTCCGCCTCCTGGGTTCACGCCA-3' | $(1.5 \times 10^8) \times 1$ |
| TATA-less 24 | 5'-TTAAATAAAGCATGCTAGATTCAAACAACTACTATGCGGCTTTAAAAAGG-3' | $(1.5 \times 10^8) \times 1$ |
| TATA-less 25 | 5'-GACTAGGCGCTGGCGGGCGGGGGTGCGCCGGAGAGTGCCCCGGCGTGTTC-3' | $(1.5 \times 10^8) \times 1$ |
| TATA-less 26 | 5'-GCTCTGGGTGTTGTACGAAAGCGCGTCTGCGGCCGCAATGTCTGCTGAGA-3' | $(1.5 \times 10^8) \times 1$ |
| TATA-less 27 | 5'-TCACTTCTGCCTTTGTTTCCTCGCTGATACGTGTTACACAGCAAGGCCAC-3' | $(1.5 \times 10^8) \times 1$ |
| TATA-less 28 | 5'-CCCCCGTACGTCACGTCGCCGCGGCCCCGCCCCCGCGCGGGCACCTCCCC-3' | $(1.5 \times 10^8) \times 1$ |
| TATA-less 29 | 5'-GACTACGTTTCCCAGGAGGCTTCGCGCGGACGCCCGGGCGGGGCTGTGCG-3' | $(1.5 \times 10^8) \times 1$ |
| TATA-less 30 | 5'-GGCGTCACAGGTCCTGACAGGGAAGAAGTTGGCAGGTCCTGGCAGGGGAC-3' | $(1.5 \times 10^8) \times 1$ |
| TATA-less 31 | 5'-GGAGAGAGGGCGCGGGAGCGAGGGCGGGCGGACGGGAGGGCCGTCAGAGC-3' | $(1.5 \times 10^8) \times 1$ |
| TATA-less 32 | 5'-AGTCCTCAGCTCACTCCCACTGCTGTCCCTGGCAGTGCAGAAGTCCAGGG-3' | $(1.5 \times 10^8) \times 1$ |
| TATA-less 33 | 5'-TATTTTTCCGGGTCATTGAAAGGAAATAAAATTCTCTGTGAAAAGTGTCT-3' | $(1.5 \times 10^8) \times 1$ |
| TATA-less 34 | 5'-TCCCTGAGCCAGACTGGATTAGGATGCCTCGCGACTAGGGGTCCAGAGAC-3' | $(1.5 \times 10^8) \times 1$ |
| TATA-less 35 | 5'-AGGAACACCCAGAGCCAGTTTTGGGAGCTGCTGGCTTCCCAAAAGCCCTT-3' | $(1.5 \times 10^8) \times 1$ |
| TATA-less 36 | 5'-AGGGACAGCCCCGCGGTTGTGGGTGTCCCGCGGCGCCGGAAACTGGGCGC-3' | $(1.5 \times 10^8) \times 1$ |
| TATA-less 37 | 5'-TTGGGTGTCTTCAGATTTCCTTGTCTTGAGGTCCTCACAATTACTCTACA-3' | $(1.5 \times 10^8) \times 1$ |
| TATA-less 38 | 5'-CGGGCCGGACAAAAGTCCCGCCTGCCCACGGCTTTTTGCCCGCCGCTCGT-3' | $(1.5 \times 10^8) \times 1$ |
| TATA-less 39 | 5'-GCTGGGACAGATTAGGGACCCCTCTTTACAGCAAGAAAGACTGCTCTGTG-3' | $(1.5 \times 10^8) \times 1$ |
| TATA-less 40 | 5'-ATGGAAAGGAACTGCAAGGGTTCCTTTGGGGTGATCAAAGAGGGAGACAC-3' | $(1.5 \times 10^8) \times 1$ |
| TATA-less 41 | 5'-CCTTTTATTTCCTCTTGTTTCAGACGGGAGCAGATATCTGTGGGTTCTTT-3' | $(1.5 \times 10^8) \times 1$ |
| TATA-less 42 | 5'-GAGCCTGGATTCGAGGGGAGGAGGGACGGGAGGAGGAGAAAGGTGGAGGA-3' | $(1.5 \times 10^8) \times 1$ |
| TATA-less 43 | 5'-GTCCCTTCCCAGGAATGAGAACCAGTGCCCGCCCCCCTCACAGCTTTCCA-3' | $(1.5 \times 10^8) \times 1$ |
| TATA-less 44 | 5'-CCCTGCTCAGATCTTTCCCATTTTCCCTCCCTTTCCCTTAGGAGCCTGTT-3' | $(1.5 \times 10^8) \times 1$ |
| TATA-less 45 | 5'-ATTAATGATTAAAATTGCCTCTACTGATTAATTATATTGATTAACTTTAT-3' | $(1.5 \times 10^8) \times 1$ |
| TATA-less 46 | 5'-TTAGCTGGGCATGGTAGCACACGTCTGTGGTCCAGCTACTTGGGAGGCTA-3' | $(1.5 \times 10^8) \times 1$ |
| TATA-less 47 | 5'-GGATGATTTAGGGACTGTTGGCTAGAACTGCCCTTGCCTACCGCCCTACT-3' | $(1.5 \times 10^8) \times 1$ |
| TATA-less 48 | 5'-TGACAATCTTCTTCCTTCCCCTGGCCACCTCTCTGCCCACTTGCTTCCTC-3' | $(1.5 \times 10^8) \times 1$ |
| TATA-less 49 | 5'-TCATCATGCTTCTACGAGGGAGATATCATTATCCCCATTTTGCAGATGGG-3' | $(1.5 \times 10^8) \times 1$ |
| TATA-less 50 | 5'-AACGTGGGGGACGGAGGCGGAAGCAGCTGGCCAAGCCGAGGTCTGTGATT-3' | $(1.5 \times 10^8) \times 1$ |

Table S1 continued...

| | | |
|---|---|---|
| coding region 1 | 5'-CAGAGGAACCCTAGAGGAGCTGTACCAGCACCCAGGTCCAGGAGGCTTGC-3' | $(1.5 \times 10^8) \times 1$ |
| coding region 2 | 5'-GATGGGGGGCGGGGGCAGGGACCATGGCAGAGCGAGAGTTCTAGGAGAGA-3' | $(1.5 \times 10^8) \times 1$ |
| coding region 3 | 5'-ACCTGGCCCCTGCAGCCTCGTCAGGCCTCTGTCCAACCTTTTCCTGTGCC-3' | $(1.5 \times 10^8) \times 1$ |
| coding region 4 | 5'-GCGCTATGGATTTCAGAGGAGTTGCAGGAATGGGGCTGCGGGGCGCACAG-3' | $(1.5 \times 10^8) \times 1$ |
| coding region 5 | 5'-CCACTCCCCTTTGCTCACATTCCCACCTGAGCTCCACCTTCTCAGATGAG-3' | $(1.5 \times 10^8) \times 1$ |
| coding region 6 | 5'-ACTAGCCTTGGGTTTTAATAGAACCTGGAAAGCTCTGTTTTGCATTTTCG-3' | $(1.5 \times 10^8) \times 1$ |
| coding region 7 | 5'-TGGCTCAATTTGAAGGTGTTTATTTTGTGTGTGCCGTGGGGCACATGGAG-3' | $(1.5 \times 10^8) \times 1$ |
| coding region 8 | 5'-CGGCCCTGCGCGCCCGGCCTCTCAGCCCCGTGCTCCCCCCAGGTGATGCC-3' | $(1.5 \times 10^8) \times 1$ |
| coding region 9 | 5'-CACTTTGGGAAGCTGAGGTGGGTGGATTGCCTGAGGTCAGGAGTTCAAGA-3' | $(1.5 \times 10^8) \times 1$ |
| coding region 10 | 5'-GAATAAAGGCAATTTGCTAACTTTCTCGCTAAATAGGATTTGGTTTCTAT-3' | $(1.5 \times 10^8) \times 1$ |
| coding region 11 | 5'-AAAATTGGCCAGGTGCAGTGGCTCACGCCTGTAATCCCAGCACTTTGGGA-3' | $(1.5 \times 10^8) \times 1$ |
| coding region 12 | 5'-TTATACAGAGCGACAAGCATCCTCTCTCAGGAAGAGCCCTCCCATCTGGG-3' | $(1.5 \times 10^8) \times 1$ |
| coding region 13 | 5'-CATTTTAATGCTTCTCTGTAATTTTACATTTCTTTTTTGTGTGTTGGGGA-3' | $(1.5 \times 10^8) \times 1$ |
| coding region 14 | 5'-CGGCGTGACCTGGGGTTCCCTGCCCCACCCCGGGCCTGTTCCACACTCTG-3' | $(1.5 \times 10^8) \times 1$ |
| coding region 15 | 5'-GGAGCGGCTGCCACTGGCGCGCCTGCTGCGCTACCTGCGGGACGACGGGC-3' | $(1.5 \times 10^8) \times 1$ |
| coding region 16 | 5'-CAAGCAAAGTCCTGCGGCCCACGCGGGAAGGCGCCCTCTTCGCGGCGCTG-3' | $(1.5 \times 10^8) \times 1$ |
| coding region 17 | 5'-GTGGGAACCAGTGCAGGGCACTCAGGAAGTCCGACCGCCCAGCGGGCCGG-3' | $(1.5 \times 10^8) \times 1$ |
| coding region 18 | 5'-TCTGATTTGCCCAATGCCGTGAGAAGATGTCGAGGAGCAGCTTCTGCTTT-3' | $(1.5 \times 10^8) \times 1$ |
| coding region 19 | 5'-ACCTCAGCCCTTCATCGACTACCAAGGCTGATCCAGCAGCCTGACCCCTC-3' | $(1.5 \times 10^8) \times 1$ |
| coding region 20 | 5'-CGGGCGGTCCTGAGCCAGGCCGTGCTCCCCACGCTCCCGATAGGTGATGC-3' | $(1.5 \times 10^8) \times 1$ |
| coding region 21 | 5'-CCTCTCCTCACAGAACTGAAGGGCCCTTGCTACCTTTAATCTGGGGGTTC-3' | $(1.5 \times 10^8) \times 1$ |
| coding region 22 | 5'-GTATTAATTATTCATTGTGTCTTTTATTACACAAATAAGGCACAGATTTT-3' | $(1.5 \times 10^8) \times 1$ |
| coding region 23 | 5'-GTTTTGTTTTTGAGACGGAGTCTCGCTCTGTCGCCCAGGCTGGAGTGCTG-3' | $(1.5 \times 10^8) \times 1$ |
| coding region 24 | 5'-GTGAGAAGCCTAGCGCCATAGTCCCTCGCCCTGCAAAGCTCAGAGGCTTC-3' | $(1.5 \times 10^8) \times 1$ |
| coding region 25 | 5'-CTCACTGGACGCTGCCTGCTGTGTCGAGGGGCCATGTGCCCAGGGCTCCT-3' | $(1.5 \times 10^8) \times 1$ |
| coding region 26 | 5'-AAGGCGGGAAATGACCCCTCCGCCGTCCCGTTACTCTCCGGGGCCAACGT-3' | $(1.5 \times 10^8) \times 1$ |
| coding region 27 | 5'-GGGCCGCCTCTCGCCGAGCACTTCAGGGGGTGTAGGGACCTGTGACCTGG-3' | $(1.5 \times 10^8) \times 1$ |
| coding region 28 | 5'-CTGCACTGGACTCAGTCTCCGGAGTGGGGCTGGGCTCCTCACCGGGCCTT-3' | $(1.5 \times 10^8) \times 1$ |
| coding region 29 | 5'-AGCATTTCTTCATATGTTTGTTGGCCACTTGTATATCTTCTTTTGAGAAG-3' | $(1.5 \times 10^8) \times 1$ |
| coding region 30 | 5'-TTTGCCTAATGGACCCAGCGCCTGGCCGCAGGCAGAGCTGCCCACCAGTC-3' | $(1.5 \times 10^8) \times 1$ |
| coding region 31 | 5'-CCCGTGTCTCTGCCCCGTCCCCCGTGTCTCTGCTCCGTCCCGTGTCTCTG-3' | $(1.5 \times 10^8) \times 1$ |
| coding region 32 | 5'-ATATGAATATTTAGCTTTGCTCCTGCTTTCTTGCTGAAGATAGGAGCTGT-3' | $(1.5 \times 10^8) \times 1$ |
| coding region 33 | 5'-TGTTCCCCGCCGCGGCGGCGCACATACCCTTCCCTAAGCTCAGGGCGTTC-3' | $(1.5 \times 10^8) \times 1$ |
| coding region 34 | 5'-TGGAGTCTCACTCTGTTGCCCAGGTTGGAGTGCAGTGGCACCATCTTAGC-3' | $(1.5 \times 10^8) \times 1$ |
| coding region 35 | 5'-TGAGCGGATCGCCTGAGGTCAGGAGTTTGAGACCAGCCTGGCCAACATGG-3' | $(1.5 \times 10^8) \times 1$ |
| coding region 36 | 5'-CCAAGGAAGCCTTACTGTGCTCGAGGTATTTGAGCCCAGCCTTTTTCCAG-3' | $(1.5 \times 10^8) \times 1$ |
| coding region 37 | 5'-CACAGGCAGGCAGCTCACAGAGTTCCCAGGCAGGTGTGCAGACAGCCGGC-3' | $(1.5 \times 10^8) \times 1$ |
| coding region 38 | 5'-CATGAAGTCTGAAGGCACTCAGACTCCTAACCCTTGTAATGCGACCTACT-3' | $(1.5 \times 10^8) \times 1$ |
| coding region 39 | 5'-GAGTGCCCTGGGCTTGGGGCGGCCGGGTCCACATGGGAGCCTCCCTCCCT-3' | $(1.5 \times 10^8) \times 1$ |
| coding region 40 | 5'-ACCTACCCATCCATTCATCCATTCCTTCATTAACCCATCCATGCACTCAT-3' | $(1.5 \times 10^8) \times 1$ |
| coding region 41 | 5'-CTTTGCAACCACTGGTCTACTTTCTCTTTTTTCTTTTTTTTTGAGACAAA-3' | $(1.5 \times 10^8) \times 1$ |
| coding region 42 | 5'-CCATGCCGGCCTGCCCTCCACTCCCGGAATCTACAAATGCATTCAAGGGC-3' | $(1.5 \times 10^8) \times 1$ |
| coding region 43 | 5'-GTATGAGGCTGTCACACTGACTCCATCAGCCCTCCTGCCTTGGCTGAAGT-3' | $(1.5 \times 10^8) \times 1$ |
| coding region 44 | 5'-AGGCTGGTCTCAAACTCCCGACCTCAGGTGATCCACACGTCTCGGCCTCC-3' | $(1.5 \times 10^8) \times 1$ |
| coding region 45 | 5'-CCTCATCTCTTACAGGTCACCAGTTAATTGTGTTGGAGGAAGCTCTGGGA-3' | $(1.5 \times 10^8) \times 1$ |
| coding region 46 | 5'-TTAGACATCTGAATCTCAGGAAACAAACAATGGAAGATAAACATCCGCAT-3' | $(1.5 \times 10^8) \times 1$ |
| coding region 47 | 5'-GTGGCCTGTTGGGATGGGGGAGAGACCCGGGGAAATATTTGGGATGACTG-3' | $(1.5 \times 10^8) \times 1$ |
| coding region 48 | 5'-TGGCTCTACAGGCAGCAGTGGCCATCAGGCAGCCAGCATCCCCCAGGGCG-3' | $(1.5 \times 10^8) \times 1$ |
| coding region 49 | 5'-GCTGCCATCTTGGCTCACTGCAACCTCCCTGCCTGATTCTCCTGCCTCAG-3' | $(1.5 \times 10^8) \times 1$ |
| coding region 50 | 5'-CCTGGACCTGAAGTCCATGGCAGACCGGGCTCGGAGCTTGGCTAAAGGAG-3' | $(1.5 \times 10^8) \times 1$ |

Table S1 continued...

| | | |
|---|---|---|
| TATA-containing Δ TATA-box 1 | 5'-GAGCATATTCTTCTATTCTTAATTTTATTATTTTTGCCTTAT-3' | $(1.5 \times 10^8) \times 1$ |
| TATA-containing Δ TATA-box 2 | 5'-AGCAATTAGCCAGGGAATGGCGTCAGGGAGACTCACTGGGCT-3' | $(1.5 \times 10^8) \times 1$ |
| TATA-containing Δ TATA-box 3 | 5'-TGATTGGCTACTTTGTTCGCATGCACGCGCGGGCGCGAGGCC-3' | $(1.5 \times 10^8) \times 1$ |
| TATA-containing Δ TATA-box 4 | 5'-TGGCGGGAGCCCTGGTGCCGGCGCGACCCGCGGACACACAGT-3' | $(1.5 \times 10^8) \times 1$ |
| TATA-containing Δ TATA-box 5 | 5'-AGAGCCTTGGTTAAAACCCCGTTCTCATCACACTGACCTGGT-3' | $(1.5 \times 10^8) \times 1$ |
| TATA-containing Δ TATA-box 6 | 5'-CAAGACCGGTATAGCTACGTTATCAATGAGAATAGGGGAGTG-3' | $(1.5 \times 10^8) \times 1$ |
| TATA-containing Δ TATA-box 7 | 5'-AGCCACGCCCAGCCGAGGGACGGCAGGTCTAGCAGACTAACC-3' | $(1.5 \times 10^8) \times 1$ |
| TATA-containing Δ TATA-box 8 | 5'-CTAGGGAGGATGTGGCGGGCCCCACCCAGGCCAGCCGGCTCT-3' | $(1.5 \times 10^8) \times 1$ |
| TATA-containing Δ TATA-box 9 | 5'-GCGGGCGGGGCCGAACGTGGGGGCGGGAGGCCAGGCTCGTGC-3' | $(1.5 \times 10^8) \times 1$ |
| TATA-containing Δ TATA-box 10 | 5'-TCTGACATGTTCTGAGCTCTCCAAAGCCCACTGCCAGTTCTC-3' | $(1.5 \times 10^8) \times 1$ |
| TATA-containing Δ TATA-box 11 | 5'-CAGGGAGGATCTGGGGCAAGGAAACTTCCAAACCTTCCAAAC-3' | $(1.5 \times 10^8) \times 1$ |
| TATA-containing Δ TATA-box 12 | 5'-GAACACCCTCGAAGTGTGGGCCAAAAAGGCAGCCTTGAATAC-3' | $(1.5 \times 10^8) \times 1$ |
| TATA-containing Δ TATA-box 13 | 5'-AAAGGAAAGGAGGAAGATGGCTTTGGGAGGAGGAGAGGGAAG-3' | $(1.5 \times 10^8) \times 1$ |
| TATA-containing Δ TATA-box 14 | 5'-TTCCTTTTAGTAGGTTCCTCATTTTACCATTTCAAACATCTT-3' | $(1.5 \times 10^8) \times 1$ |
| TATA-containing Δ TATA-box 15 | 5'-CACATATTCAGCTCCAGCACCTCATCTGCTCTGACTTCCCCA-3' | $(1.5 \times 10^8) \times 1$ |
| TATA-containing Δ TATA-box 16 | 5'-TGGAACAGAGACTGTATGAGGCAAAAGAAGTCTTCTGGGAAT-3' | $(1.5 \times 10^8) \times 1$ |
| TATA-containing Δ TATA-box 17 | 5'-GCTAGGACTCGACCTGGGCGCCTCCAGGGAAACGCTAAGGGG-3' | $(1.5 \times 10^8) \times 1$ |
| TATA-containing Δ TATA-box 18 | 5'-AATATTTGCGGGCCATCCCAGGCTGCTGTCAGAATATAACAG-3' | $(1.5 \times 10^8) \times 1$ |
| TATA-containing Δ TATA-box 19 | 5'-TTCCTGTTGCCGGGACGCACGTGATGAGCGCACGGGCTGCGG-3' | $(1.5 \times 10^8) \times 1$ |
| TATA-containing Δ TATA-box 20 | 5'-TAAAGATACATATGTATGCATTTGTGTGTGTATGTATGTGTA-3' | $(1.5 \times 10^8) \times 1$ |
| TATA-containing Δ TATA-box 21 | 5'-CTGTGGTGAGAATTAATCTATGATGTGTTAGACTCTCCAGCT-3' | $(1.5 \times 10^8) \times 1$ |
| TATA-containing Δ TATA-box 22 | 5'-CTATGTTATTTTAATTTTTACTATTGTGTGTGTGTGTGTGTA-3' | $(1.5 \times 10^8) \times 1$ |
| TATA-containing Δ TATA-box 23 | 5'-CTTTAAAACAGAAGAAAAATAACTACCTTAAATTTAAAATTG-3' | $(1.5 \times 10^8) \times 1$ |
| TATA-containing Δ TATA-box 24 | 5'-AGGAGCCACATCTTTCTGTGGCCTATGAAAAACAGAGTGCTT-3' | $(1.5 \times 10^8) \times 1$ |
| TATA-containing Δ TATA-box 25 | 5'-CTCTTTGGACGGGATGTACTATAAATTGCTATGTCAAGCAGT-3' | $(1.5 \times 10^8) \times 1$ |
| TATA-containing Δ TATA-box 26 | 5'-AAAAAACTTCCCTGGGGGCTCTCTCCCTGGCTAAAGATAGTA-3' | $(1.5 \times 10^8) \times 1$ |
| TATA-containing Δ TATA-box 27 | 5'-TGGGCGGGGTCCAGCGGCGGGCGGAAGGCGGGGCGTGGGGGT-3' | $(1.5 \times 10^8) \times 1$ |
| TATA-containing Δ TATA-box 28 | 5'-CAGTGAATGCCACACTGTTCCTCAGCCTTTCCTAGGGGTGTG-3' | $(1.5 \times 10^8) \times 1$ |
| TATA-containing Δ TATA-box 29 | 5'-GGGAGGGCCGAGTAGGCGACGGTGAGGTCGACGCCGGCCAAG-3' | $(1.5 \times 10^8) \times 1$ |
| TATA-containing Δ TATA-box 30 | 5'-AGTTAAAACCAACATAACATTTCTAGCAAATAATGGGAAGAA-3' | $(1.5 \times 10^8) \times 1$ |
| TATA-containing Δ TATA-box 31 | 5'-CGCCCGAGGCCATCGCCACCTGCTGCCACTAGCCAAGCCGCG-3' | $(1.5 \times 10^8) \times 1$ |
| TATA-containing Δ TATA-box 32 | 5'-AGTTCCTGGGAGATACTGGACGGCCAAAGCATCTTCTGAGAC-3' | $(1.5 \times 10^8) \times 1$ |
| TATA-containing Δ TATA-box 33 | 5'-AAGGGGTGTATAACTGAGACCTCAGAGAGAAAACTCACCACC-3' | $(1.5 \times 10^8) \times 1$ |
| TATA-containing Δ TATA-box 34 | 5'-GGCCCGGGACCAGCGCCGCGGCGGCTGCGGCGAGGCCGGCAG-3' | $(1.5 \times 10^8) \times 1$ |
| TATA-containing Δ TATA-box 35 | 5'-ATAAAGAACCACCATATGCTAACTAACCCTCCTGATTTTATC-3' | $(1.5 \times 10^8) \times 1$ |
| TATA-containing Δ TATA-box 36 | 5'-ATGGTCTGATATGCCCTATAAAATGACATATAAGGCACCCTA-3' | $(1.5 \times 10^8) \times 1$ |
| TATA-containing Δ TATA-box 37 | 5'-AGCATATCATCAAGAAATAACTAGGCAACCAGCAGCTCCCGG-3' | $(1.5 \times 10^8) \times 1$ |
| TATA-containing Δ TATA-box 38 | 5'-CGGGCGGGGCCGAACGTGGGGGCGGGAGGCCAGGCTCGTGCC-3' | $(1.5 \times 10^8) \times 1$ |
| TATA-containing Δ TATA-box 39 | 5'-ACATGCCACAAAGGCACAGCGGTGGGAATCAGAGCACTTCAG-3' | $(1.5 \times 10^8) \times 1$ |
| TATA-containing Δ TATA-box 40 | 5'-AAAAGGCCCTTCGGAGTATTTATAATAATTATGATTATTATT-3' | $(1.5 \times 10^8) \times 1$ |
| TATA-containing Δ TATA-box 41 | 5'-GTCTGTCTGTCTATCAGAGTTTCCAAATGAACTGTAGTTTGT-3' | $(1.5 \times 10^8) \times 1$ |
| TATA-containing Δ TATA-box 42 | 5'-AGCTCTCTTCATTTTATAACTTTCAGGAGGATCTCATGTATC-3' | $(1.5 \times 10^8) \times 1$ |
| TATA-containing Δ TATA-box 43 | 5'-ACGTCACCCCGGGCTTGGATGTGATCAGGGAGCTGGGGAGAA-3' | $(1.5 \times 10^8) \times 1$ |
| TATA-containing Δ TATA-box 44 | 5'-AGCCTGACCAACATGGTGAAACCCCGTCTCTACTAAAAATTA-3' | $(1.5 \times 10^8) \times 1$ |
| TATA-containing Δ TATA-box 45 | 5'-CTAAGCCTCATCCATCACTGATCAGTCTTCACGTAAGGATTT-3' | $(1.5 \times 10^8) \times 1$ |
| TATA-containing Δ TATA-box 46 | 5'-GCAACCGCTCACAGTCTGCGCTCCTGGTACACGCGCTTCAAC-3' | $(1.5 \times 10^8) \times 1$ |
| TATA-containing Δ TATA-box 47 | 5'-CCCTGAAGCCCGCAGGTCCCCGCCTCGGCAGCCACGGGACAC-3' | $(1.5 \times 10^8) \times 1$ |
| TATA-containing Δ TATA-box 48 | 5'-TGTGTGTGTATATTTTGAGATGAAATCTCACTCTTTTGCCCA-3' | $(1.5 \times 10^8) \times 1$ |
| TATA-containing Δ TATA-box 49 | 5'-GGGTGTCTCAACATCCTTCATGGTGGCCACTAGTAACGCAGG-3' | $(1.5 \times 10^8) \times 1$ |
| TATA-containing Δ TATA-box 50 | 5'-ACGTCGAGCTTCAAAAATTTTTGCAATTTTTAGGTTTTATAT-3' | $(1.5 \times 10^8) \times 1$ |

### 3 Supplementary Figures

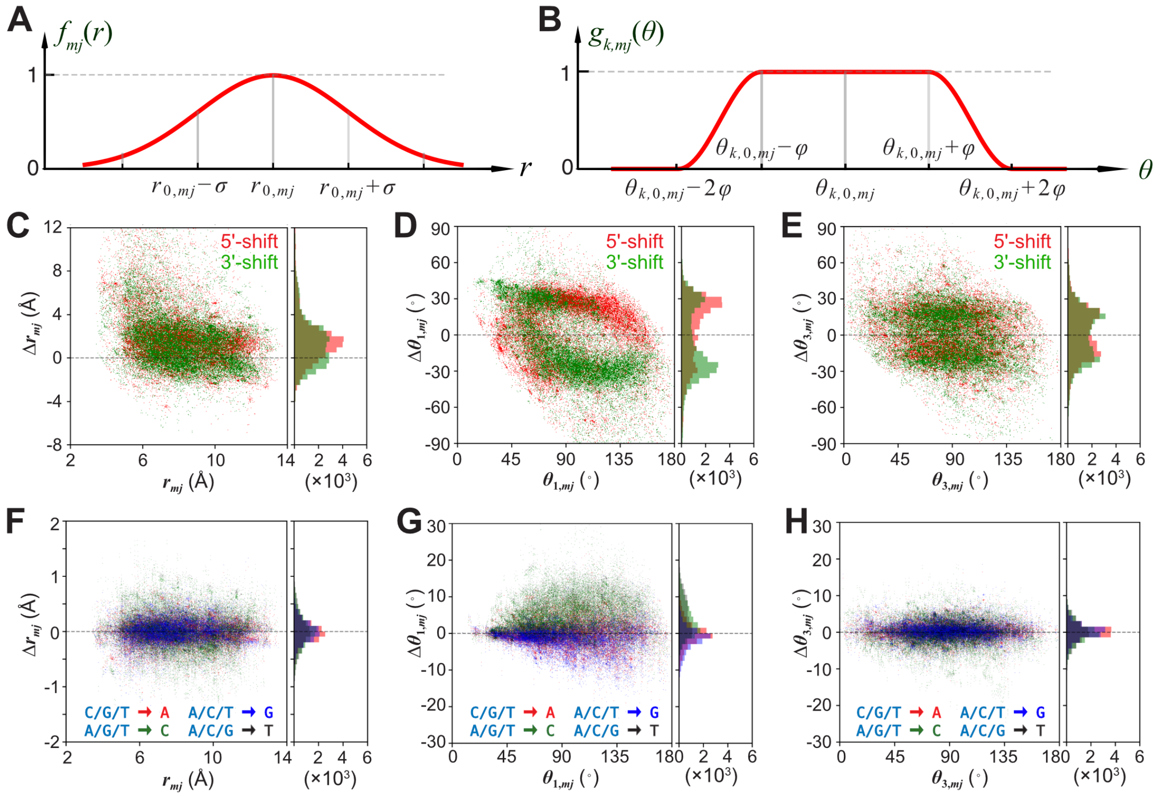


Figure S1: Graphs of the modulating functions and determination of parameters ($\sigma$ and $\varphi$). (A) and (B) show the two modulating functions, $f_{mj}(r)$ and $g_{k,mj}(\theta),\ k = 1, 2, 3$, respectively. Roughly speaking, $\sigma$ and $\varphi$ control the "widths" of the modulating functions. (C-E) show the distributions of $r_{mj}$, $\theta_{1,mj}$, and $\theta_{3,mj}$ as well as their changes ($\Delta r_{mj}$, $\Delta\theta_{1,mj}$, and $\Delta\theta_{3,mj}$) under the 1-bp shift tests (see Fig. 3A for definition), respectively. (F-H) are about $r_{mj}$, $\theta_{1,mj}$, $\theta_{3,mj}$ and their changes under the mutation tests ( Fig. 3C).

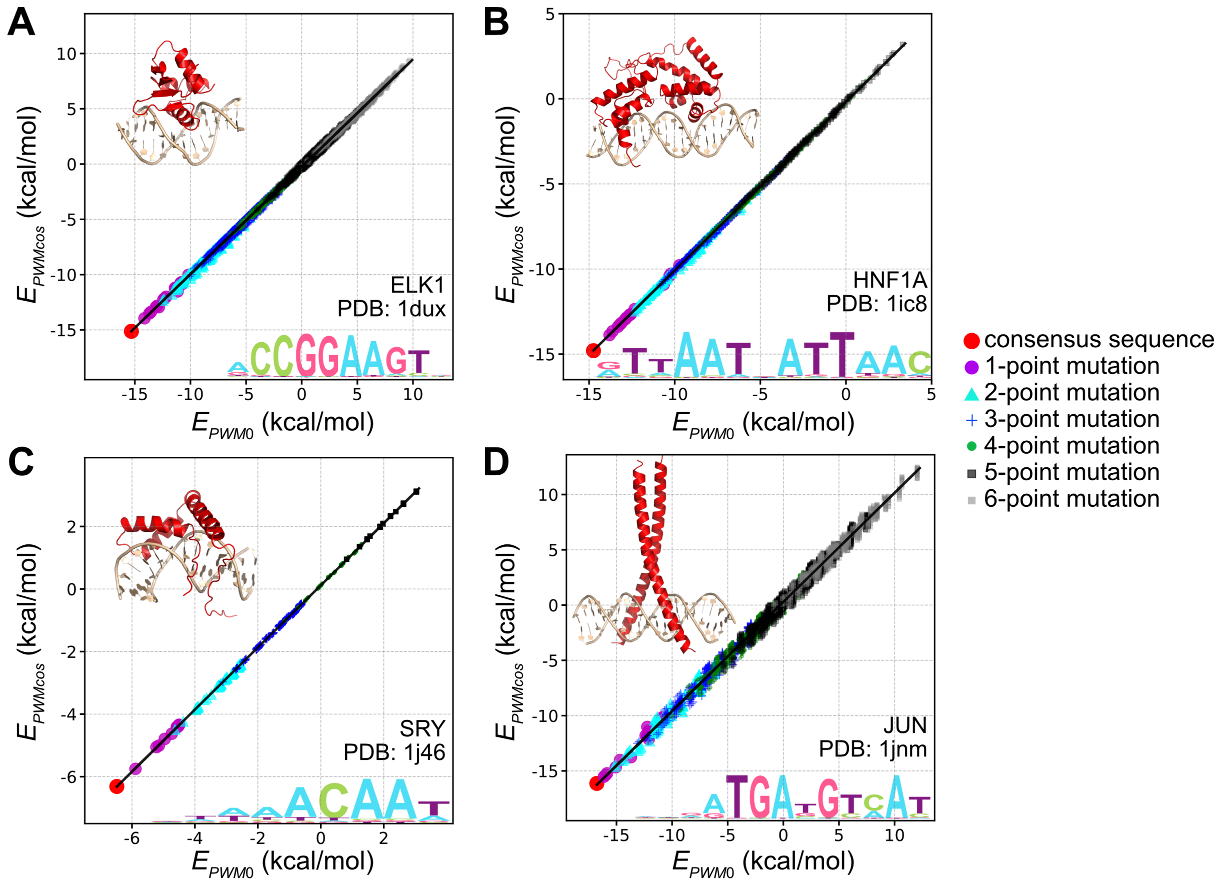


Figure S2: Correlation between the energy directly computed with PWM ($E_{PWM}$) and the energy calculated with our new model ($E_{PWMcos}$) for four proteins, ELK1 (A), HNF1A (B), SRY (C), and JUN (D), when they bind to the consensus sequences or the mutated sequences. Color and shape of dots indicate different number of mutations introduced, as shown by the legends on the right side. For each protein, we show the PDB structures and the sequence logos in the insets. Note that due to limited computational power, for long consensus sequences (such as for HNF1A and JUN) we did not introduce all possible mutations, but instead randomly generated a large number of mutated sequences for the test.

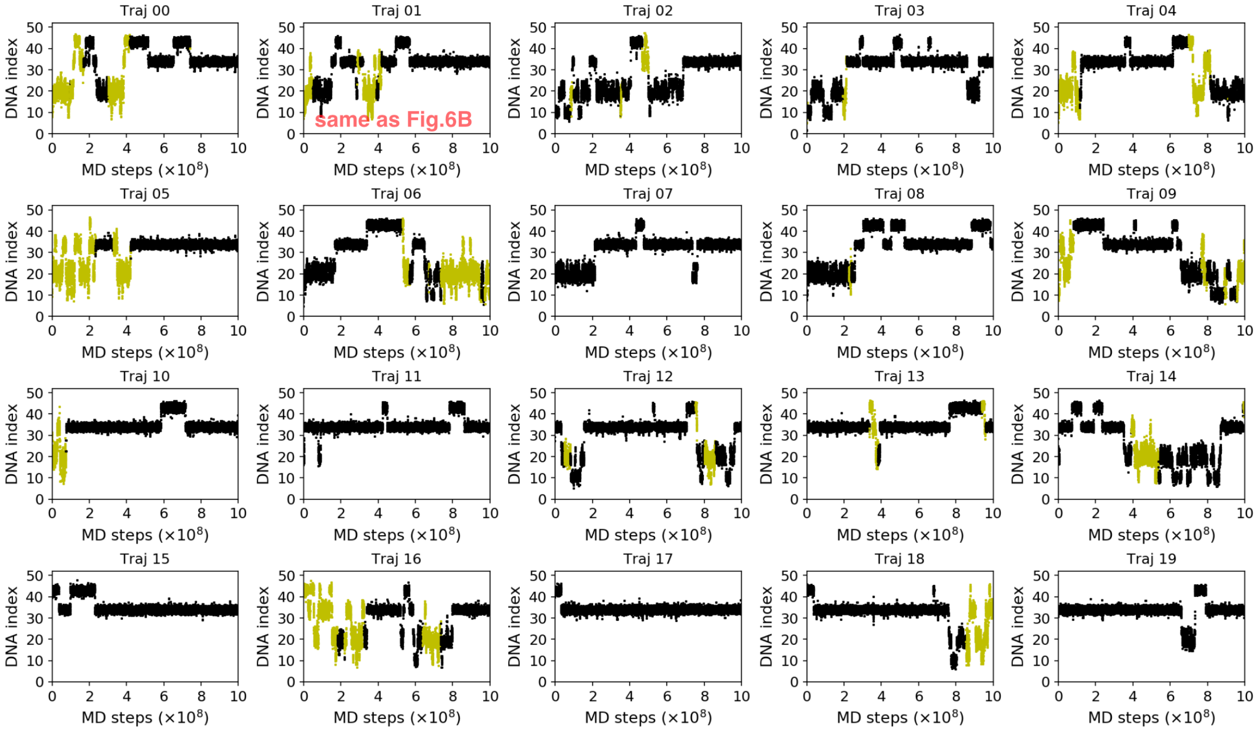


Figure S3: Time series of PU.1 binding positions on DNA in 20 independent $10^9$-step MD trajectories. The color of the dot represents the relative orientation of PU.1 on DNA (see Fig. 6A and 6B).

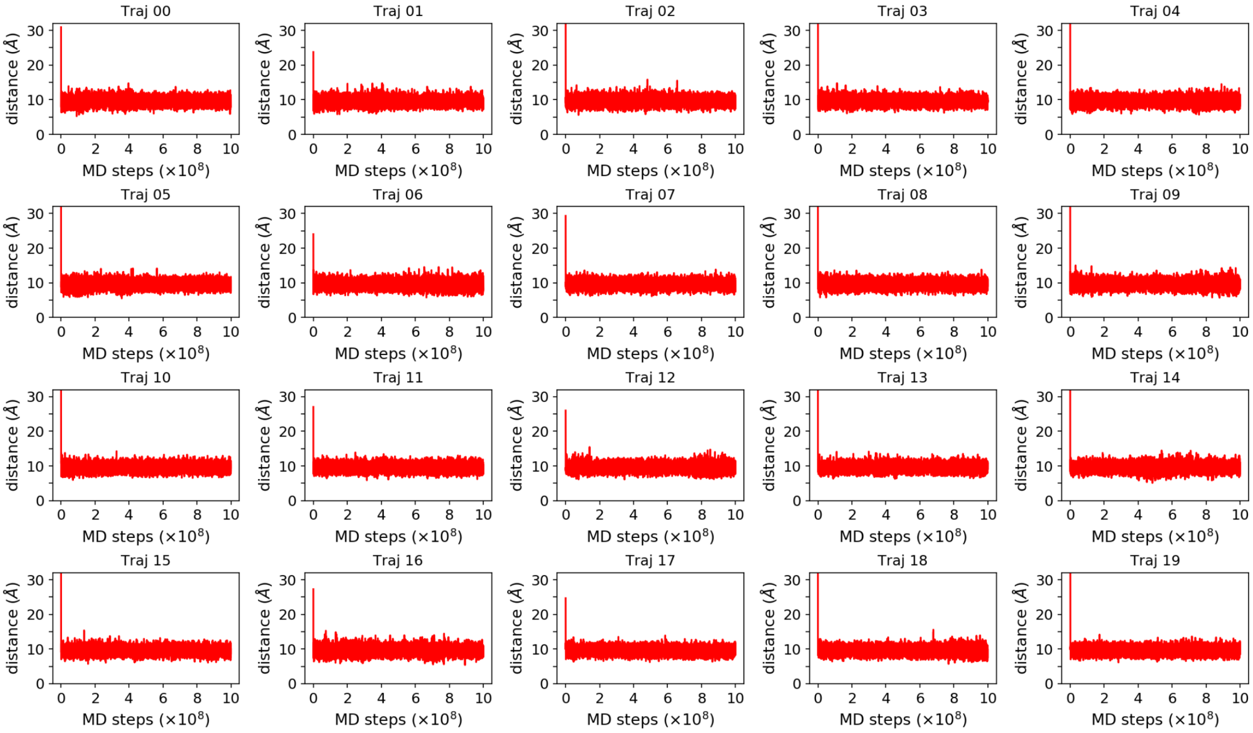


Figure S4: Time series of the distance from center of mass of PU.1 to the nearest DNA particle.

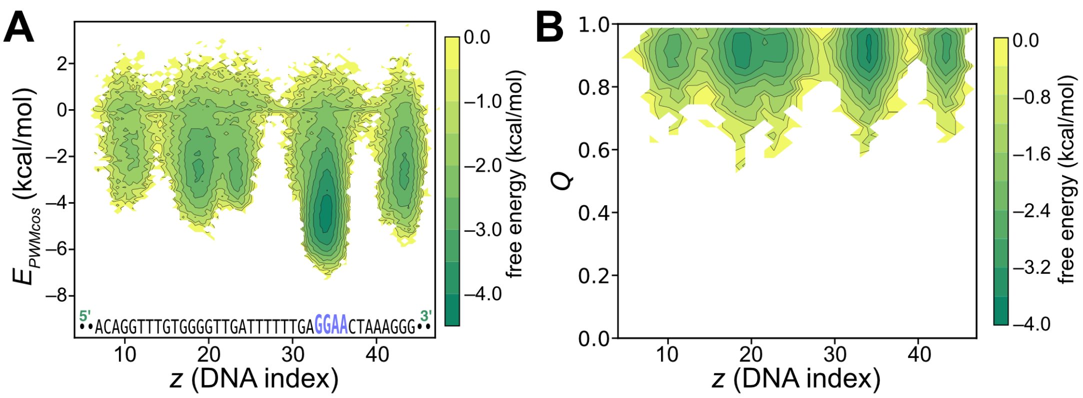


Figure S5: Free energy surface projected onto two-dimensional profiles of $E_{PWMcos}$-$z$ (A) and $Q$-$z$ (B). $Q$ represents the native-ness of DNA-binding interface of PU.1, as defined in the main text.

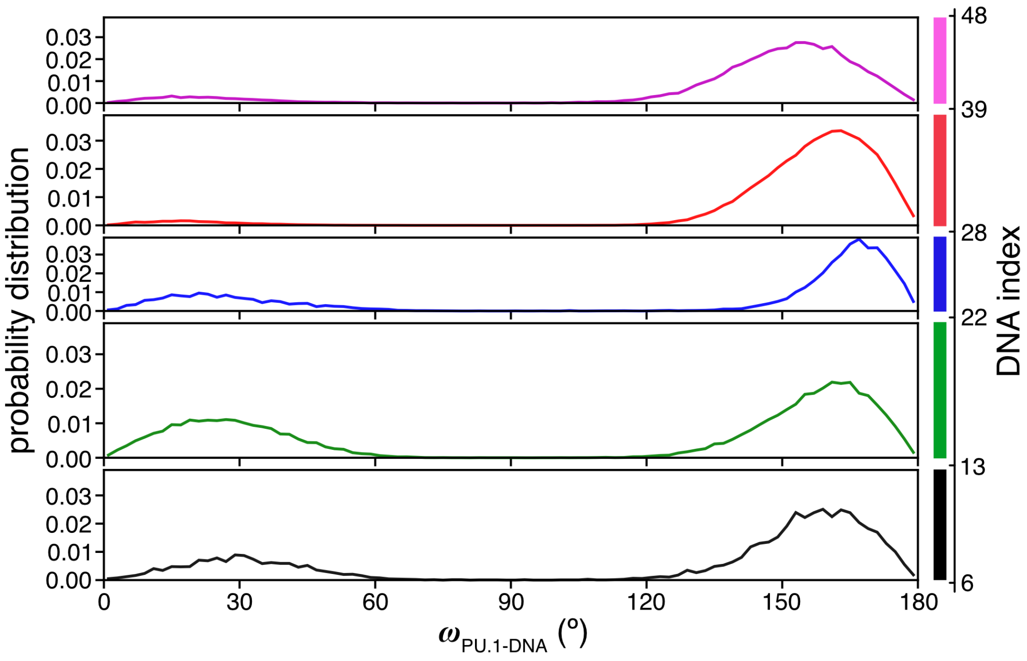


Figure S6: Probability distributions of $\omega_{\text{PU.1-DNA}}$ when PU.1 was bound to different regions of DNA. The division of DNA sequences into pieces (regions) are based on the 1D-FES ($F_1$) plotted in Fig. 6C. Each region here corresponds to a local minimum in the 1D-FES.

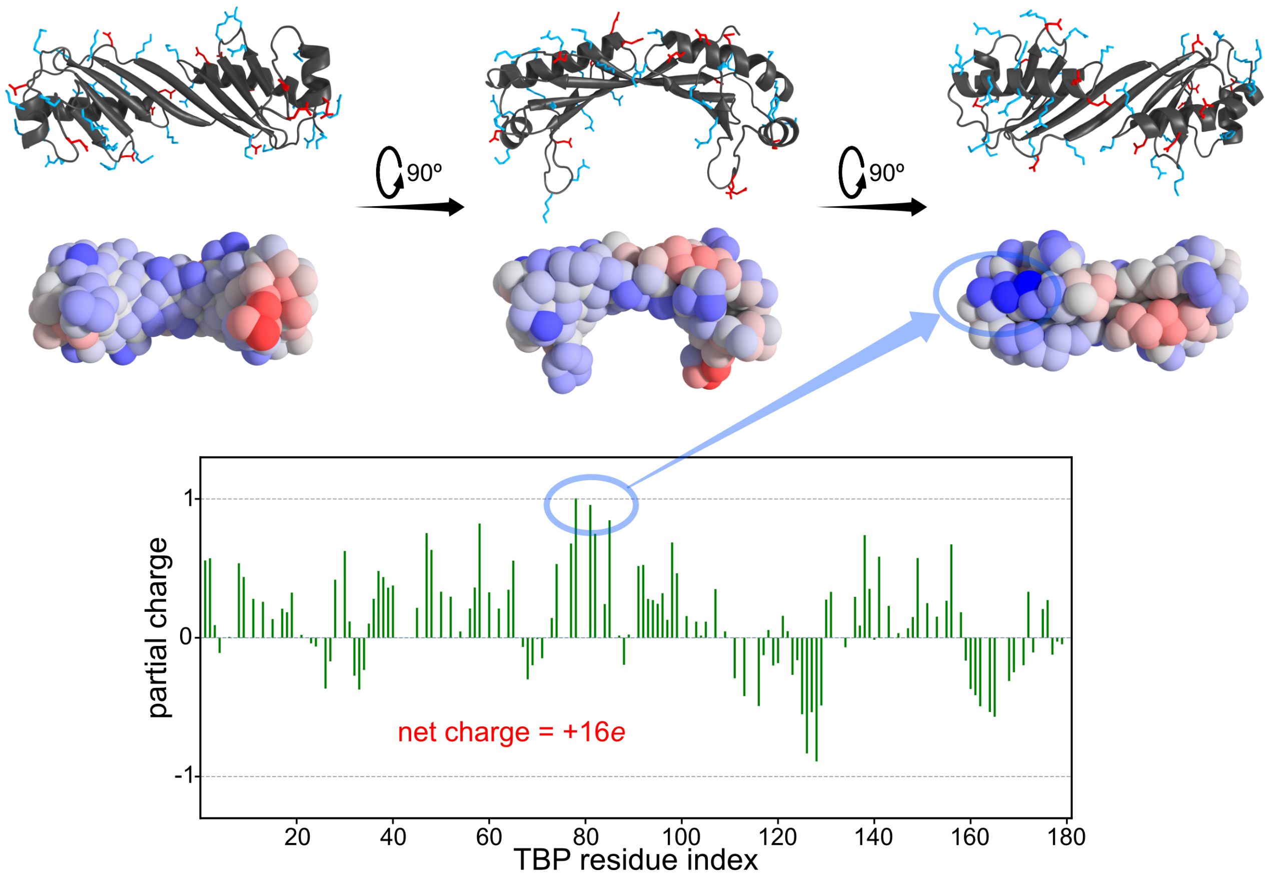


Figure S7: Distribution of charged residues on the surface of TBP. (Upper) In the first row, the atomistic structures are shown in the cartoon representation, with positively charged residues (Arg, Lys, and His, in blue) and negatively charged residues (Asp and Glu, in red) shown as sticks. In the second row, the coarse-grained (CG) structures are shown as spheres, with the color representing partial charge distributed on each CG particle, ranging from $-1.0e$ (red) to $+1.0e$ (blue). The partial charges were calculated with the RESPAC method[9]. (Lower) Bar plot of the RESPAC partial charges on TBP residue. The most positively charged residues are marked with blue circles and connected by a blue arrow to show their locations in the protein structure.

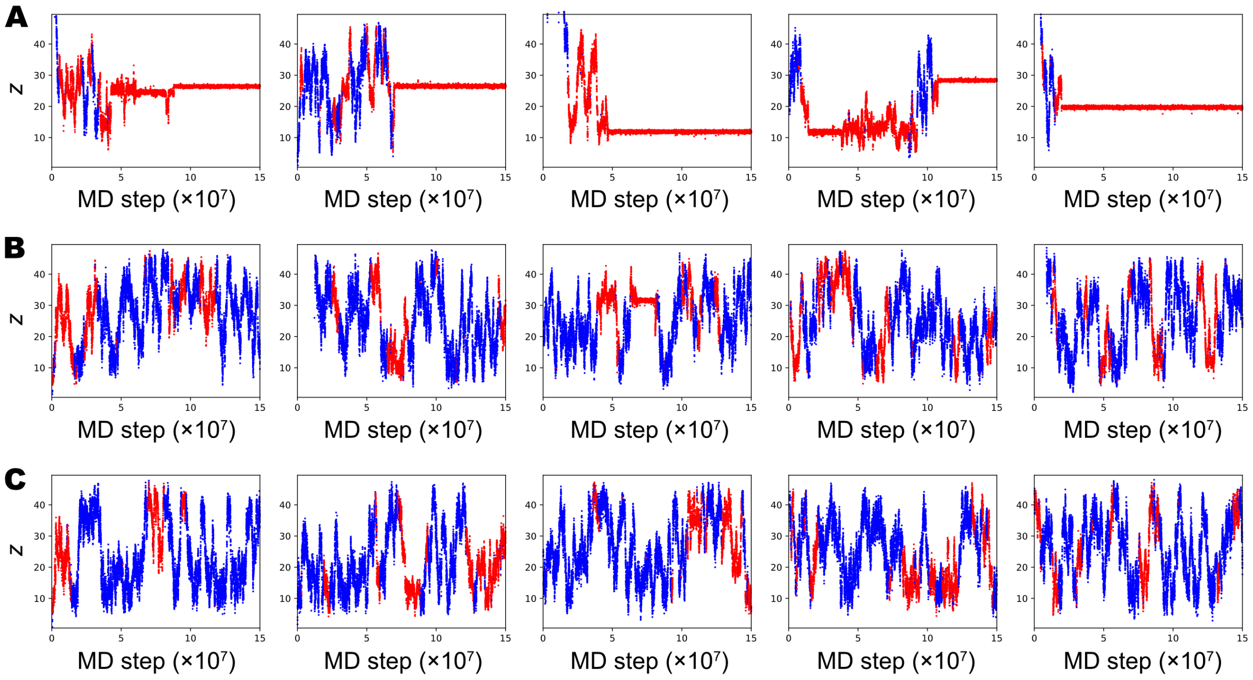


Figure S8: Representative trajectories of TBP binding to TATA-containing promoters (A), TATA-less promoters (B), and coding regions (C).

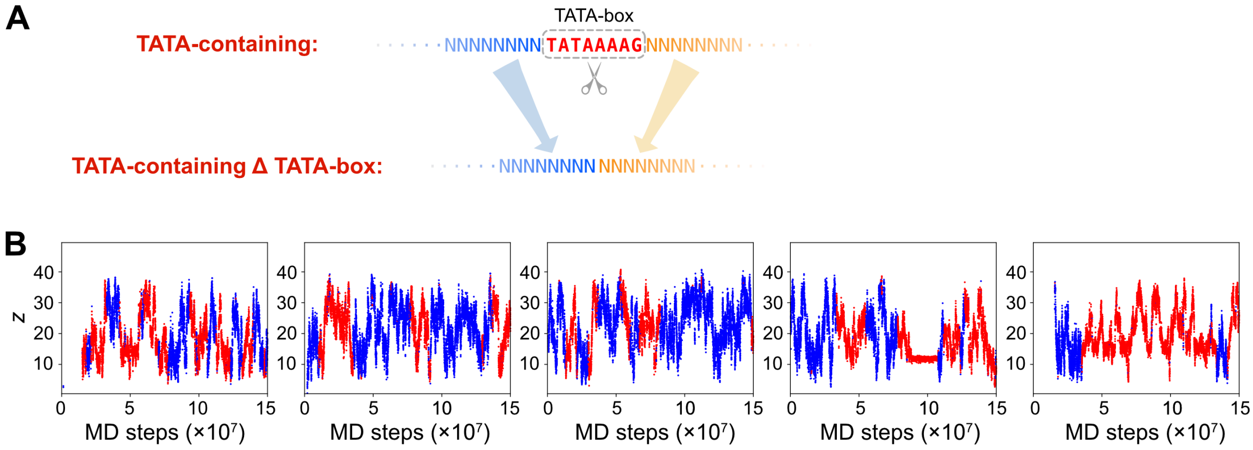


Figure S9: Simulations of TBP binding to TATA-containing promoter sequences removing the TATA-boxes (termed "TATA-containing ∆ TATA-box"). (A) Simulation DNA sequence design. We determined TATA-box sequence elements by the regular expression "TATA[AT]A[AT][AG]". The TATA-boxes were then deleted from the TATA-containing sequences and the remaining parts were used for TBP binding simulations. (B) Representative trajectories of TBP binding to the "TATA-containing ∆ TATA-box" sequences.

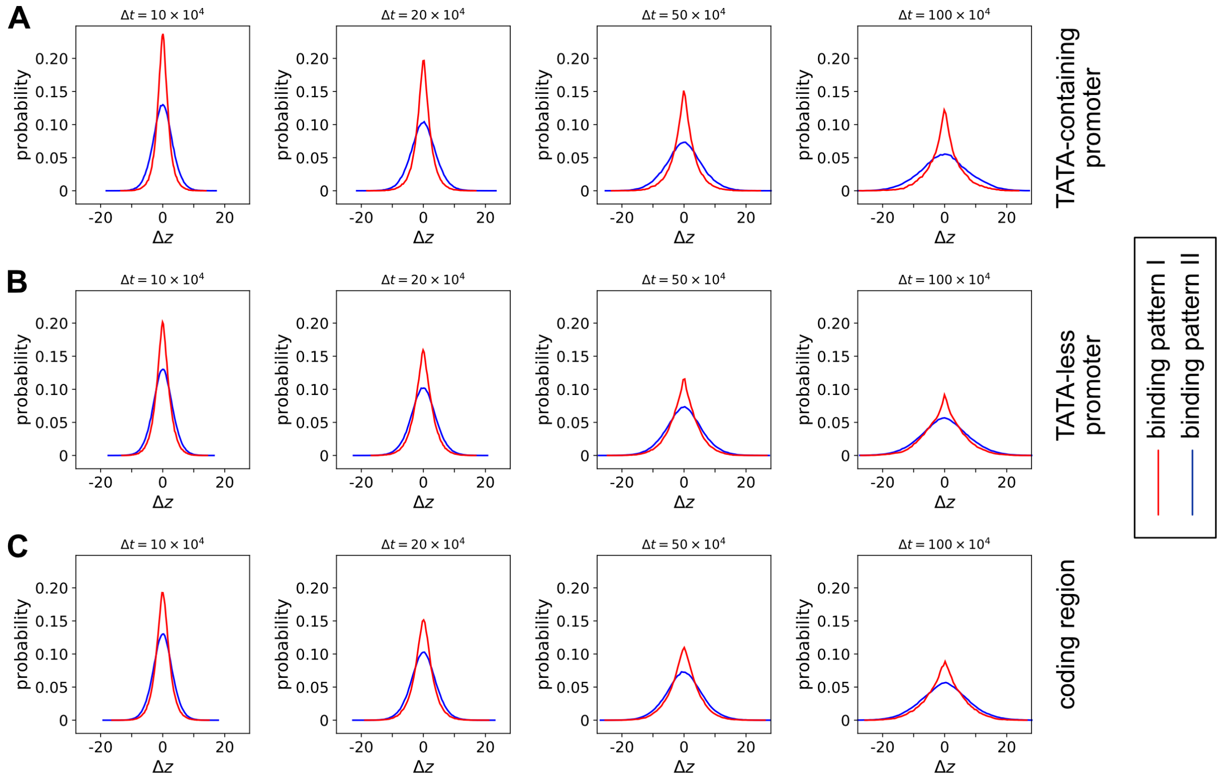


Figure S10: Time-dependent ($\Delta t$, in the unit of MD step) distributions of TBP sliding distance $\Delta z$ in simulations of TBP binding to TATA-containing promoter (A) (excluding the TATA-box specific binding frames), TATA-less promoter (B) and coding region sequences (C). Colors of curves represent different TBP binding patterns. Note that due to space limitation, we only show $\Delta z$ distributions for the first four $\Delta t$s.

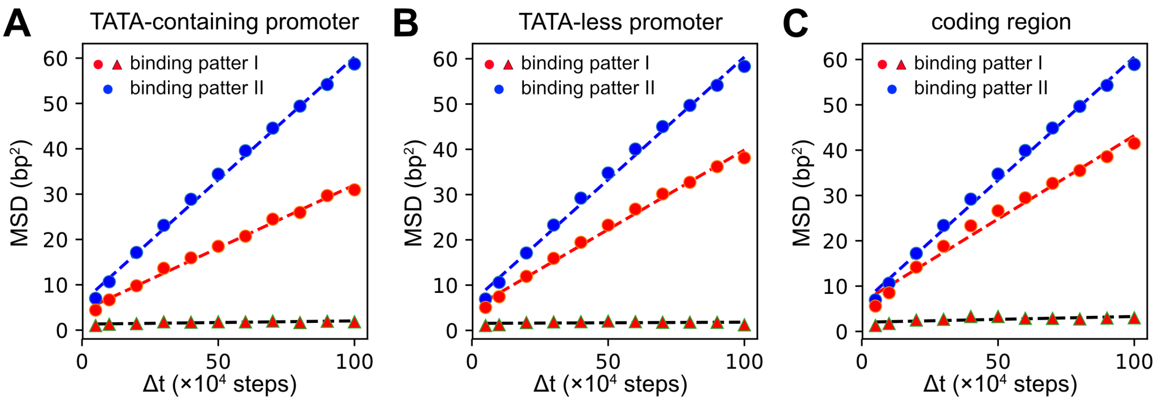


Figure S11: Fitted one-dimensional diffusion coefficient ($D_1$) of TBP on TATA-containing promoter (A), TATA-less promoter (B) and coding region sequences (C). For the binding pattern I, we first performed a double-Gaussian fit to the $\Delta z$ distributions (Fig. S10) and then calculated $D_1$ for each decomposed Gaussian (red triangles and red circles). Whereas for binding pattern II, single Gaussian fitted well with the $\Delta z$ distributions and we calculated $D_1$ based on MSD of $\Delta z$ as a function of $\Delta t$ (see Supplementary Methods).

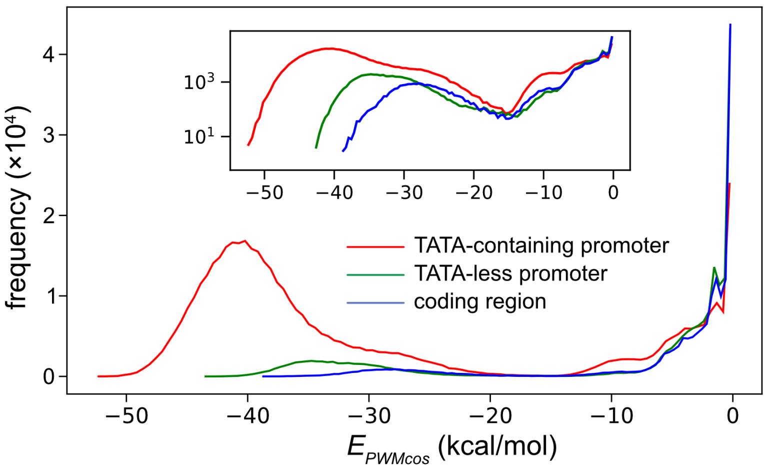


Figure S12: Distribution of $E_{PWMcos}$ in simulations of TBP binding to TATA-containing promoter (red), TATA-less promoter (green), and coding region (blue) DNA sequences. Inset is the same but with the vertical axis in $\log_{10}$ scale.
